# Allele-specific expression reveals multiple paths to highland adaptation in maize

**DOI:** 10.1101/2022.07.15.500250

**Authors:** Haixiao Hu, Taylor Crow, Saghi Nojoomi, Aimee J. Schulz, Matthew B. Hufford, Sherry Flint-Garcia, Ruairidh Sawers, Ruben Rellan-Alvarez, Juan M. Estévez-Palmas, Jeffrey Ross-Ibarra, Daniel E. Runcie

## Abstract

Maize is a staple food of smallholder farmers living in highland regions up to 4,000 meters above sea level worldwide. Mexican and South American highlands are two major highland maize growing regions, and population genetic data suggests the maize’s adaptation to these regions occurred largely independently, providing a case study for parallel evolution. To better understand the mechanistic basis of highland adaptation, we crossed maize landraces from 108 highland and lowland sites of Mexico and South America with the inbred line B73 to produce F_1_ hybrids and grew them in both highland and lowland sites in Mexico. We identified thousands of genes with divergent expression between highland and lowland populations. Hundreds of these genes show patterns of convergent evolution between Mexico and South America. To dissect the genetic architecture of the divergent gene expression, we developed a novel allele-specific expression analysis pipeline to detect genes with divergent functional *cis-*regulatory variation between highland and lowland populations. We identified hundreds of genes with divergent *cis-*regulation between highland and lowland landrace alleles, with 20 in common between regions, further suggesting convergence in the genes underlying highland adaptation. Further analyses suggest multiple mechanisms contribute to this convergence. Our findings reveal a complex genetic architecture of *cis*-regulatory alleles underlying adaptation to highlands in maize. Although the vast majority of evolutionary changes associated with highland adaptation were region-specific, our findings highlight an important role for convergence at the gene expression and gene regulation levels as well.

## Introduction

Highland maize is cultivated in cold, mountainous regions worldwide, at altitudes of up to 4,000 meters above sea level (masl) and with mean growing season temperatures below 20°C (Lothrop 1994; Hartkamp et al. 2000)⍰. The International Maize and Wheat Improvement Center (CIMMYT) estimates that more than 6 million hectares (Mha) are used for highland maize production worldwide, mainly in developing countries where it is grown by smallholder farmers as one of the main sources of calories in their diet (Lothrop 1994; Zambrano et al. 2021)□. Mexico (~2.9 Mha, 46.6%) and South America (~0.6 Mha, 9.4%) are two major highland maize producing regions and are geographically separated from each other. Highland maize landraces (open-pollinated traditional varieties) in central Mexico and South America have distinct morphological characteristics from lowland tropical or temperate germplasm (Janzen et al. 2022)□, including purple stems, drooping leathery leaves, weak roots, tassels with few branches, conical-shaped ears (Anderson and Cutler 1942), and a changed biochemical response to UV radiation (Casati and Walbot 2005). They also have other specific characteristics that make them suitable to live in high-elevation climates, including frost tolerance and improved seedling emergence, growth, and grain filling at low temperatures (Eagles and Lothrop 1994)□.

These consistent differences between highland and lowland landraces indicate that highland maize has undergone considerable local adaptation since its introduction to highland environments in the past 6200 years (Piperno and Flannery 2001)⍰. However, we still know little about the genetic basis of highland adaptation in maize: What genes were involved? Was adaptation driven by standing genetic variation or novel alleles? Is the genetic basis of adaptation parallel between populations from different geographic regions? Recent population genetic studies have begun to paint a complex and divergent picture of highland adaptation between Mexican and South American maize. Genome-wide SNP data shows strong population structure in maize landraces from Mesoamerica and South America (Van Heerwaarden et al. 2011;⍰ Takuno et al. 2015). Several studies using population genetic data (Hufford et al. 2013; Pyhäjärvi et al. 2013; Calfee et al. 2021; Rodríguez-Zapata et al. 2021)⍰ identified genomic loci that were introgressed from a wild ancestor of maize, *Zea mays ssp. mexicana* (hereafter *mexicana*) found exclusively in the highlands of central and northern Mexico (De Jesús Sánchez González et al. 2018), suggesting that alleles contributing to highland adaptation may have been acquired by crossing with pre-adapted relatives. Three of these loci have been well characterized: *Inv4m* (Hufford et al. 2013; Crow et al. 2020⍰), *mhl1* (Hufford et al. 2013; Calfee et al. 2021) and *HPC1* (Rodríguez-Zapata et al. 2021), and the *mexicana* alleles are found almost exclusively in landraces from the Mexican highlands. Wang et al. (2017)⍰ found no evidence for substantial spread of *mexicana* haplotypes to South America and Takuno et al. (2015)⍰ found < 1.8% of SNPs and 2.1% of genes showing evidence for convergent evolution between Mesoamerican and South American highland populations. However, in a recent genome-wide scan with high-density SNPs, Wang et al. (2021) ⍰identified 10,199-11,345 SNPs and 1,651-2,015 genes with evidence for population divergence between highland and counterpart lowland populations in Central America and South America, respectively, including 10.7% of SNPs, 15.0% of genes, and flowering time pathway showing evidence of parallel adaptation between Andes and Mexican highland landrace populations. The extent of parallelism in adaptation to highlands is important because it can indicate whether alleles beneficial for highland adaptation in one geographic region are likely to also be beneficial in another or whether adaptation is likely constrained by a limited set of possible loci or if multiple different adaptive paths are available (Lee and Coop, 2017; Wang et al. 2021).

Population genetic scans using SNP markers can efficiently discover loci that have diverged between populations, indicating a potential role in adaptation. However, discovering mechanisms controlled by these loci remains a challenge. While predicting the function of protein-coding variants is possible, we have little ability to predict the function of non-coding variants, including those affecting gene regulation. For example, although the 13 Mb *Inv4m* locus has been known about for more than a decade (Hufford et al. 2013)□ and appears to play a role in flowering time (Romero Navarro et al. 2017)□, the mechanisms underlying its role remained unclear. Gene expression analysis can provide a link between sequence variation and molecular mechanisms, particularly by discovering expression patterns of groups of genes that share common biological functions or attributes (Maleki et al. 2020). Crow et al. (2020) developed two populations segregating for highland and lowland alleles at this locus and measured gene expression effects of the locus across nine tissues. They identified 39-607 genes per tissue that were consistently regulated by *Inv4m* in both families, and gene set enrichment analyses suggested a role of the locus in the regulation of photosynthesis and several other biological processes. Other studies have begun to use gene expression to study the process of highland adaptation in maize as well. Kost et al. (2017)□ measured expression variation among landraces from three distinct elevational zones (highland, midland and lowland) and identified two co-expression modules correlated with temperature-related environmental parameters. Rocío Aguilar-Rangel et al. (2017)□ used allele-specific expression to study *cis*-regulatory divergence between the highland landrace Palomero Toluqueño and the modern inbred B73 and identified 2,386 genes with divergent expression caused by the different genotypes. These expression studies are limited however, in their ability to describe the complexity and genetic architecture of gene regulatory adaptation at the population level where evolution occurs.

In this study, we used population-level allele specific expression (ASE) analyses to identify gene expression traits that have diverged between highland and lowland populations of Mexican and South American maize landraces. We selected maize landraces from 108 highland and lowland sites that cover broad growing regions of highland maize in Mexico and South America and crossed them with a common inbred line B73 to produce F_1_ hybrids (F_1_s). We planted the F_1_ families at two locations that represented highland and lowland environments in Mexico. Our primary objectives were to (i) identify genes that show evidence for adaptive divergence in *cis-*regulation between high and low elevation landraces in Mexico and South America; (ii) identify candidate gene pathways and functional groups that underwent directional selection for gene regulation during adaptation to highland climates; and (iii) gain insights into the convergent evolutionary patterns of highland adaptation between populations in Mexico and South America. We first identified genes with divergent expression between highland and lowland populations. We then differentiated the two alleles of each gene using ASE and identified genes with divergent *cis-*regulation between highland and lowland alleles. To achieve the population-level ASE analysis, we developed a novel analysis pipeline that can accurately measure the ASE of each individual at the gene level using RNAseq data alone. We discovered hundreds of genes with divergent *cis*-regulation between highland and lowland landrace alleles in the Mexican and South American populations, respectively. Of these, 20 genes were in common between populations, suggesting a low level of convergence at the gene regulation level underlies highland adaptation in maize.

## New Approaches

Allelic read counts are the starting point of all ASE analyses (Castel et al. 2015)□. Most ASE analyses have been done either based on individual SNPs (Shao et al. 2019; Zhou et al. 2019; Li et al. 2021)⍰ or by integrating allelic read counts across SNPs within a gene (Lemmon et al. 2014, Fan et al. 2020)□. Gene-level ASE ratios are more robust because they are based on more total reads, and in a population sample SNP-level ASE ratios cannot reliably be compared across individuals because many SNPs are individual-specific. Therefore, most existing studies using ASE have been based on a single F_1_ individual (Rocío Aguilar-Rangel et al. 2017; Shao et al. 2019; Zhou et al. 2019, but see Lemmon et al. 2014 who used 29 F_1_s from different maize and teosinte parents to study the genetics of maize domestication)□, so the generality of the discoveries to whole populations was unclear.

We have developed a novel analysis pipeline that can accurately measure ASE of each individual at the gene level using RNAseq data alone, and efficiently detect genes with common functional variation in *cis*-regulatory regions that have diverged between populations. First, we crossed maize landraces from 108 highland and lowland sites in Mexico and South America with a common inbred line B73 to produce F_1_ hybrids and took advantage of this genetic design to phase heterozygous SNPs of each F_1_ sample based on the B73 reference genome. Then, we extracted reads that were assigned to either of the two parental origins at all overlapping loci with heterozygous SNPs into separate BAM files and counted the reads overlapping each gene feature in each BAM file. These gene counts are the allelic expressions of the maternal and paternal alleles of each gene, respectively. Finally, we tested for *cis*-regulatory divergence between highland and lowland populations in the Mexican and South American populations by analyzing the average difference in landrace allele-specific expression (relative to B73 allele-specific expression) between F_1_s derived from highland and lowland landraces.

Our methodology can efficiently detect genes showing *cis*-regulatory divergence between populations. In addition, gene-level ASE ratios estimated with our method can be used to identify gene-trait relationships relevant to hybrid breeding through transcriptome-wide association studies (TWAS). In such programs, candidate lines are evaluated by crossing to common testers. TWAS using ASE can pinpoint causal gene regulatory traits underlying key performance traits of interest, enabling further targeted gene editing for genetic improvement.

## Results

### Geographical origins and population structure of maize landraces

We selected 108 maize landraces from CIMMYT’s germplasm bank representing highland and lowland sites (one landrace accession per site) across broad geographical regions of Mexico and South America where maize landraces are cultivated (fig. 1A; supplementary table S1). Individuals from highland (> 2000 masl) and lowland (<1000 masl) sites were paired latitudinally (within 1 degree latitude) and chosen such that all pairwise distances were greater than 50 km (fig. 1A).

**Fig.1.**
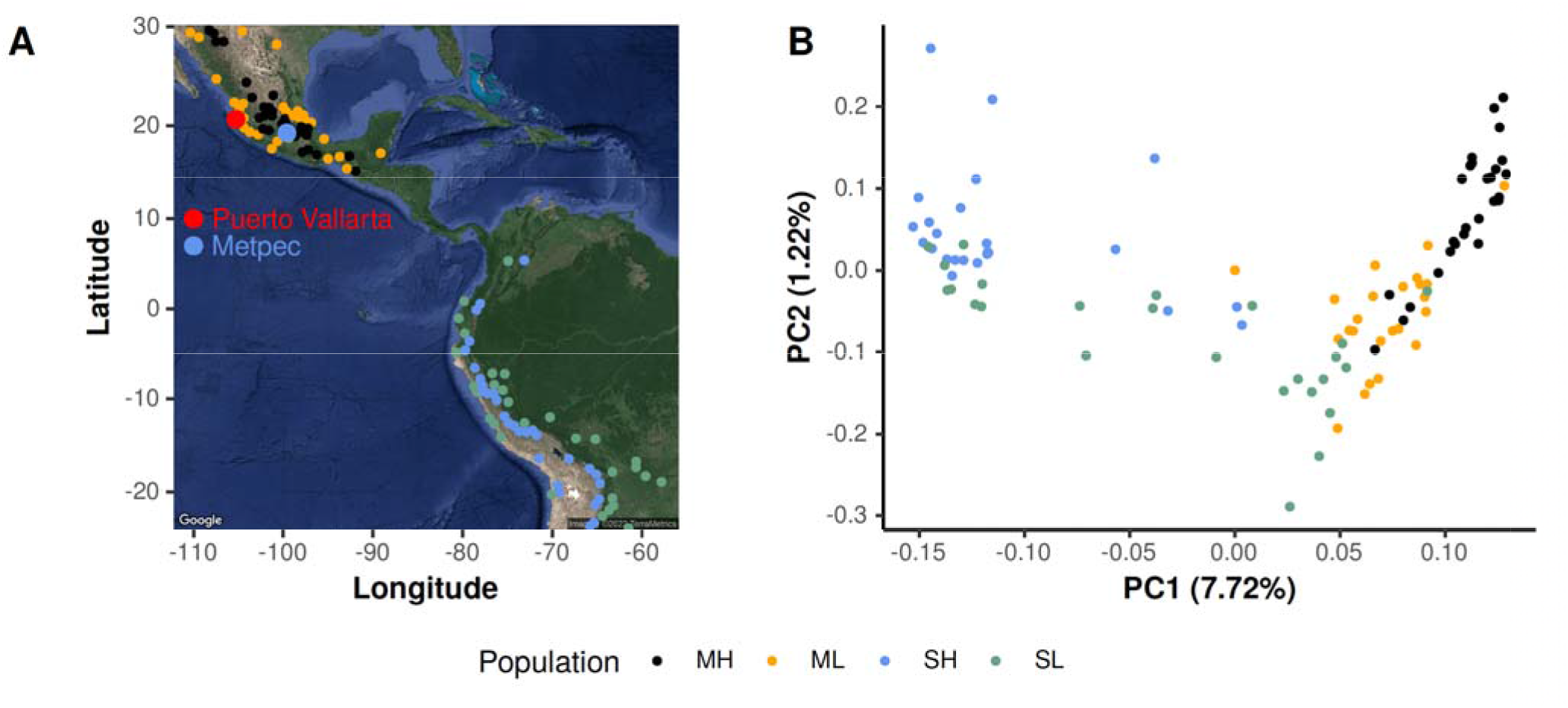
The Geographical origins (A) and genomic relationships (B) of the 108 maize landraces used as paternal parents of the F1 populations. MH=Mexican Highland, ML=Mexican Lowland, SH=South American Highland, and SL=South American Lowland. In Figure A, the larger dots represent physical positions of the two field trials, and the smaller dots represent physical positions where the 108 maize landraces were collected.

We did whole-genome skim sequencing of a single plant of each of these 108 landraces and performed a principal component analysis (PCA, fig. 1B) to study the genetic structure of the landraces. The first two principal components (PCs) separated the landraces into four populations (Mexican Highland, Mexican Lowland, South American Highland, South American Lowland). The genomic relationships of the 108 maize landraces estimated here were consistent with Janzen et al. (2022)□ who used a different individual from each of the same landrace populations genotyped with DArTseq-Based SNP markers (Wenzl et al. 2004)□. Our results were also consistent with patterns of genetic structure reported by Van Heerwaarden et al. (2011)□ using a small SNP panel of 1,127 accessions of maize landraces.

### Highland and lowland landraces show widespread divergences in gene expression

We measured gene expression in F_1_ hybrids derived from the 108 landraces described above in two leaf-derived tissues sampled from two locations: leaf tip and leaf base samples from a fully expanded leaf of a V4 plant from each F_1_ family in each of two field blocks at the highland site in Metepec, Mexico at 2620 masl, and leaf tip samples from a comparably staged leaf from a single plant from each F_1_ family in a single field block at the lowland site in Puerto Vallarta, Mexico at 7 masl. These tissues (hereafter site:tissues) are labeled MetLeaftip, MetLeafbase, and PvLeaftip below. In each of these three site:tissues, we tested for differences in the expression of each expressed gene between highland and lowland-derived F_1_s separately for the Mexican and South American populations, accounting for sampling effects due to time of collection and collection team, and leveraging shared signals across site:tissues using multi-variate adaptive shrinkage (mash) (Urbut et al. 2019). In total, we discovered 4,432 and 1,816 (supplementary tables S2, S3) genes with differential expression between highland and lowland derived F_1_ plants from the Mexican and South American continents, respectively, using a 5% local false sign rate (*lfsŕ*) threshold for declaring significance. Breaking these lists down by site:tissue, we discovered 1278, 3716, and 319 genes with divergent highland expression in the Mexican F_1_ families in MetLeaftip, MetLeafbase, and PvLeaftip, and 715, 1626, and 368 genes with divergent highland expression in the South American F_1_ families (fig. 2A, supplementary fig. S1). We detected many more genes with differential expression between highland and lowland landraces on each continent than between the Mexican and South American populations on average (total of 124 genes, supplementary table S3), or that were associated with latitude on either continent (total of 60 and 131 genes in the Mexican and South American populations, respectively, supplementary table S3). However, many more genes showed significant changes in expression during the approximately 1.5hr sampling window within each site:tissue (a total of 18,844 out of the 21,599 genes assayed across the 3 site:tissues, supplementary table S3, supplementary fig. S2), suggesting that the transcriptome-wide consequences of elevation adaptation were smaller than diel expression variation during the course of a morning.

**Fig.2.**
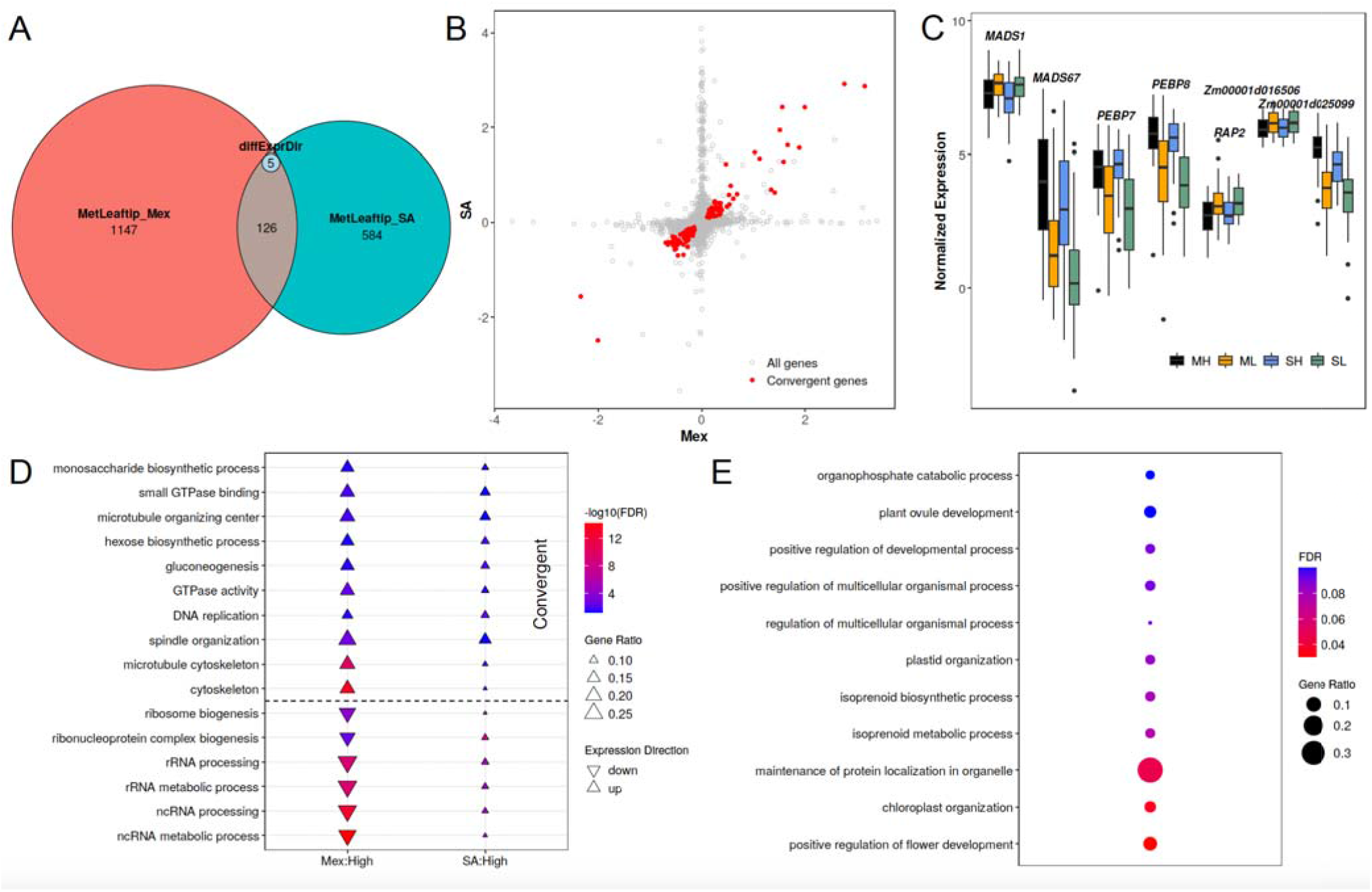
Results of gene expression analyses. (A) Numbers of differentially expressed genes between highland and lowland populations from Mexico and South America and common genes detected in both continents in the MetLeaftip tissue. The small inset in the overlapping region shows genes significant in both populations, but with opposite directions of expression change (B) Correlation of Posterior Mean highland effects between Mexican and South American population for all genes measured for gene expression (in gray) and a subset of genes showing evidence of convergent evolution (in red) in the MetLeaftip tissue. (C) Expression of flowering-related genes in the Mexican Highland (ML), Mexican Lowland (ML), South American Highland (SA), and South American Lowland (SL) populations in the MetLeaftip tissue. These flowering-related genes are identified by looking for overlapping between the convergent genes and maize flowering time candidate genes aggregated by Li et al. (2016) and Swarts et al. (2016). (D) False discovery rate (FDR) of 16 Gene Ontology (GO) terms that are significant in both Mexican and South American populations across three site:tissue. The size of each triangle indicates the enrichment ratio of this GO term, defined as ratio of number of differentially expressed genes in a GO category divided by the size of the category. We tested up-regulated and down-regulated differentially expressed genes separately and triangles and upside-down triangles represent up-regulated and down-regulated GO categories, respectively. (E) GO categorical enrichments of the genes individually classified as having convergent expression evolution in MetLeaftip and MetLeafbase.

Among these genes with differential expression in highland populations, a small minority were significantly associated with elevation in the F_1_ families of both continents. 131, 429 and 30 were detected in both continents per site:tissue, representing 18%, 26% and 8% of the lesser of the number of significant genes from either continent (fig. 2A, supplementary fig. S1, supplementary table S4). However, despite being a relatively small overlap, this is many more than expected by chance (p=2.74×10^-25^, 1.12×10^-17^, and 3.28×10^-12^ per site:tissue, respectively), and if we relax the significance threshold, the overlap percentage grows considerably larger. Furthermore, of the genes with significant responses to elevation on both continents, both the direction and magnitude of expression difference between highland and lowland populations was highly correlated (fig. 2B, supplementary fig. S3). While the estimated highland effects were positively correlated for all genes (r=0.22, 0.26, and 0.20), the effects of genes with significant effects in both populations were much higher (r=0.96, 0.94, and 0.97). We thus considered the 126, 411 and 30 genes exhibiting identical directional change of expression as having convergent evolution of gene expression between the two continents.

Because previous studies of highland adaptation in maize have described earlier flowering as a characteristic of highland landraces (Romero Navarro et al. 2017; Wang et al. 2021; Janzen et al. 2022)□, we inspected a list of maize of 886 flowering time genes and candidates aggregated by Li et al. (2016)□ and Swarts et al. (2016)□. Of these, 17 showed convergent expression differences in F_1_ families from both continents (fig. 2C, supplementary fig. S4, table 1), including four well-known transcription factors and *ZCN8* that contributes to early flowering during highland adaptation (Guo et al. 2018). Additionally, phosphatidylglycerols have been linked to the regulation of flowering through the sequestration of florigen in phloem cells (Susila et al. 2021), and we found 31 and 12 differentially expressed genes (supplementary table S5) labeled with the Gene Ontology term “phosphatidylglycerol biosynthetic process” (GO:0006655) associated with elevation from the Mexican and South American continents, respectively, using a 5% *lfsr* to declare differentially expressed genes. All of these differentially expressed genes were down-regulated in the highlands in both populations, consistent with earlier flowering. If we relax the significance threshold, for example *lfsr*=0.2, the differentially expressed genes mapped to GO:0006655 and down-regulated in the highlands in both populations grow to 50 and 36 with 30 in common (supplementary table S5). These results further support Wang et al.’s (2021) finding of convergent evolution of flowering regulation along elevational gradients in Mexico and South America.

**Table 1.** 17 flowering-related genes that showed convergent expression differences between highland and lowland-derived F_1_ families from Mexican and South American populations

| GeneID | Gene Name | Chr | Position(bp) | Description | Expression changes in highland genotypes | References |
| --- | --- | --- | --- | --- | --- | --- |
| Zm00001d022088 | MADS67 | 7 | 169,844,061 | MADS-transcription factor 67 | up | Li et al. 2016 |
| Zm00001d010752 | PEBP8/ZCN8 | 8 | 126,880,531 | phosphatidylethanolamine-binding protein8 | up | Swarts et al. 2016 |
| Zm00001d038725 | PEBP7/ZCN7 | 6 | 163,368,049 | phosphatidylethanolamine-binding protein7 | up | Swarts et al. 2016 |
| Zm00001d010987 | RAP2 | 8 | 136,009,216 | rap2 - rap2.7 orthologue (transcription factor) | down | Swarts et al. 2016 |
| Zm00001d025099 |  | 10 | 103,947,429 |  | up | Li et al. 2016 |
| Zm00001d016506 | cl27878_1 | 5 | 165,302,124 |  | down | Li et al. 2016 |
| Zm00001d048474 | MADS1/ZMM5 | 9 | 156,960,598 | transcription factor | down | Swarts et al. 2016 |
| Zm00001d049543 | CCA1 | 4 | 34,070,590 |  | down | Swarts et al. 2016 |
| Zm00001d051951 |  | 4 | 175,147,743 |  | down | Li et al. 2016 |
| Zm00001d014990 | RUP1 | 5 | 71,267,717 | repressor of UV-B photomorphogenesis homolog1 | down | Li et al. 2016 |
| Zm00001d015293 |  | 5 | 82,992,330 |  | up | Li et al. 2016 |
| Zm00001d005814 |  | 2 | 189,518,235 |  | down | Li et al. 2016 |
| Zm00001d040323 | CAL2 | 3 | 38,197,170 | calmodulin2 | up | Li et al. 2016 |
| Zm00001d022558 |  | 7 | 180,004,346 |  | up | Li et al. 2016 |
| Zm00001d023833 |  | 10 | 23,764,459 |  | down | Li et al. 2016 |
| Zm00001d046935 |  | 9 | 111,766,412 |  | down | Li et al. 2016 |
| Zm00001d012119 | JMJ11 | 8 | 168,442,999 | JUMONJI-transcription factor 11 | up | Li et al. 2016 |

Position(bp) represents starting physical position of a gene (bp; B73 AGPv4)

Beyond flowering regulation, the long lists of differentially expressed genes (supplementary table S2) themselves are difficult to parse for insights into highland elevation. Therefore, to summarize these results, we tested for enrichment of Gene Ontology categories (Wimalanathan et al. 2018) □and KEGG (Kanehisa et al. 2021) and CornCyc (Hawkins et al. 2021) □pathways among the lists of significant genes, measuring enrichment separately for up-regulated and down-regulated highland genes in each site:tissue. A total of 763 GO categories, 38 KEGG pathways and 3 CornCyc pathways were significantly enriched in at least one site:tissue at a 5% false discovery rate (FDR) (supplementary table S6). The most significant GO terms were thylakoid (GO:0009579), plastid envelope (GO:0009526), chloroplast envelope (GO:0009941).

Of these functional GO categories, 16 were identified in F_1_ families from both continents, and 10 of them were similarly enriched with up-regulated or down-regulated genes on both continents suggesting that the evolutionary changes were convergent (fig. 2D). Confirming the results above, categorical enrichments of the genes individually declared to show convergent expression evolution identified 6 and 15 terms in MetLeaftip and MetLeafbase (fig. 2E, supplementary table S7), respectively, including the terms positive regulation of flower development (GO:0009911) and chloroplast organization (GO:0009658), and also including endoplasmic reticulum (ER) retention sequence binding (GO:0046923).

To explore whether the gene expression changes could be partially explained by alterations in cell-type compositions of leaf tissues, we used a set of marker genes for 7 cell populations identified by single cell sequencing of a maize leaf (Bezrutczyk et al. 2021)□ to estimate relative cell population sizes in each sample. The first two principal components of our cell population scores clearly separated the three site:tissues (fig. 3A), and the scores explained significantly more variation among samples than expected from random subsets of genes (fig. 3B), suggesting that these gene sets captured meaningful variation, even if the precise identities of the cell populations are not clear. The first principal component of the cell population scores of the MetLeafbase sample were also unevenly distributed across the field, suggesting spatial variation in leaf anatomy or developmental stage. However, within each range of the field the highland and lowland samples from the Mexican population were clearly differentiated, and highland and lowland samples from the South American population were also clearly differentiated across 3/5 of the field (fig. 3C), suggesting that there were consistent anatomical differences between highland and lowland leaves. These anatomical differences likely cause the appearance of differential expression because different cell populations express genes at different levels.

**Fig.3.**
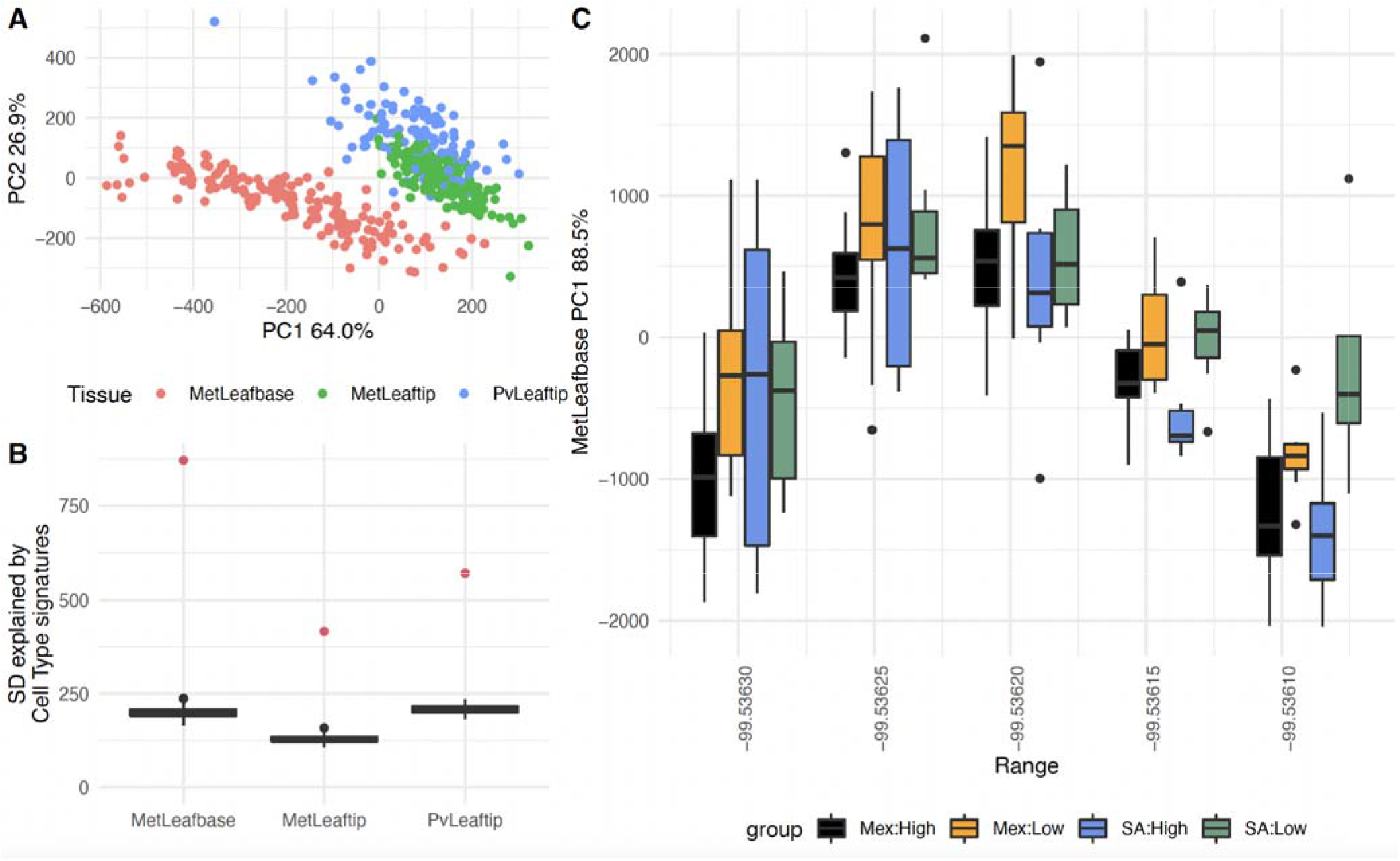
Cell type proportion inference. (A) Each point represents a single RNA sample, colored by the site:tissue and positioned according to its coordinates on the first two principal component axes of the projections onto seven sets of cell-type specific genes identified by Bezrutczyk et al. (2021)⍰ in maize leaves. (B) Red points show the standard deviation of the cell-type projection scores within each tissue. Black box-plots show the distribution of 200 randomized projection scores based on random sets of genes. (C) Distributions of the PC1 coordinates for the MetLeafbase samples, separated by population and range of the field.

We attempted to control for these anatomical differences when testing for differential expression between highland and lowland accessions by including the cell population scores as covariates. In these models the number of differentially expressed genes and enriched GO terms dropped significantly (a total of 648 genes and 0 GO terms were significant for elevation in the Mexican population, and a total of 1182 genes and 68 GO terms were significant for elevation in the South American population, supplementary table S8) suggesting that anatomical differences were the primary driver of expression differences observed above, at least for the Mexican population. However, the differential expression of flowering-related genes remained significant even after accounting for these anatomical differences.

### Development of a novel allele specific expression analysis pipeline to identify genetic loci underlying morphological and/or transcriptomic differences between highland and lowland landraces

The gene expression analysis results above point to a diverse set of expression traits associated with highland adaptation in Mexican and South American landraces; however, the genetic architecture of these differences remains unclear. While differential gene expression analyses can detect differences in thousands of expression traits, it remains possible that a small number of genetic loci might be responsible for most of these changes (Crow et al. 2020)□. On the other hand, differences in expression between the two allelic copies of each gene in each F_1_ individual can only be caused by differences in the local *cis*-regulatory region around each gene (Sun and Hu 2013)□. Therefore, we used ASE (defined as the ratio of landrace allelic count to B73 allelic count) to scan the genome for genes that have undergone divergence in the *cis*-control of gene expression between highland and lowland landraces.

To resolve major challenges (supplementary text) for ASE detection across individuals at the gene level when only RNAseq data is available, we took advantage of our genetic design involving the 108 F_1_ hybrids all crossed to the same tester line B73 (fig. 4). We developed a novel analysis pipeline for directly counting allelic reads at the gene level in each F_1_ individual. Briefly, our pipeline included three parts: First, we identified a set of high-confidence SNPs between any of the landrace parents and B73 from our low-coverage whole-genome sequencing data. Next, we used the RNAseq data to genotype and phase these SNPs within each F_1_ sample. Finally, we counted the number of reads confidently assigned to either the B73 reference or landrace genome, accounting for allelic mapping bias using the WASP algorithm (Van De Geijn et al. 2015)□. Full details are available in the Methods.

**Fig.4.**
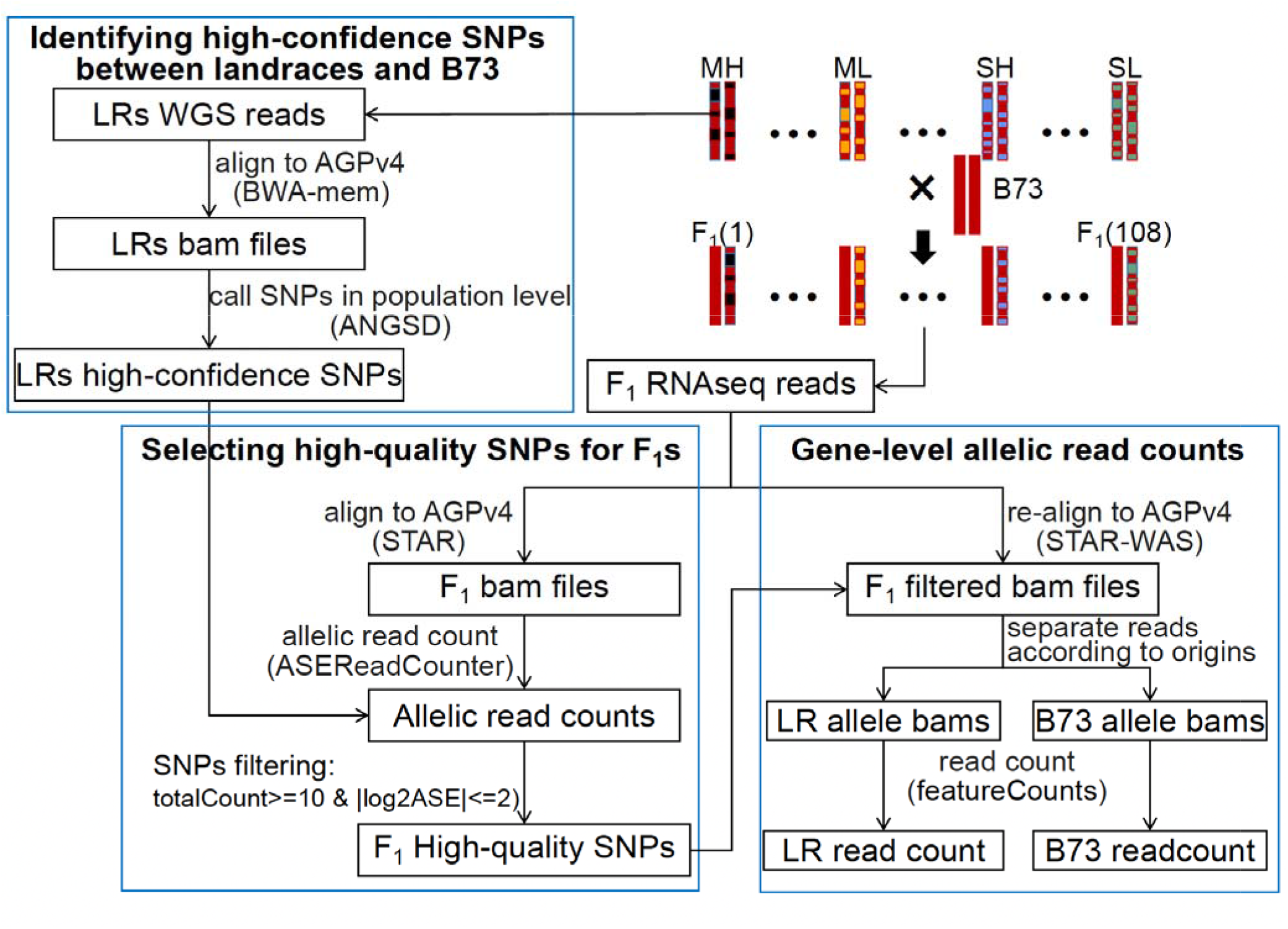
The analysis pipeline for gene-level allelic read count. LR=landrace, WGS=whole genome sequencing, AGPv4=B73 reference genome version 4, bams=bam files, MH=Mexican Highland, ML=Mexican Lowland, SH=South American Highland, and SL=South American Lowland.

To assess the reliability of our pipeline, we performed three validation analyses. First, the distribution of log2ASE ratios across all genes was approximately symmetric around zero for each sample, suggesting that we did not have strong reference bias towards the B73 allele (supplementary fig. S5A). In contrast, less stringent filtering of SNPs led to strong reference allele bias (supplementary fig. S5B). Second, the ASE values from our real data had much more variation than expected by counting variance alone, suggesting the observed variation is due to biology (supplementary fig. S6). Finally, the correlation of ASE between samples collected from two different individuals from the same F_1_ family was high for genes in genomic regions where the two individuals shared the same haplotype but much lower for genes in genomic regions where the two individuals did not share the same haplotype (supplementary fig. S7). Full details are available in the supplemental results.

### Detection of differential *cis*-regulation of landrace alleles between highland and lowland landrace populations

We tested for *cis*-regulatory divergence at the population level between highland and lowland alleles in the Mexican and South American populations by comparing ASE ratios among samples for each gene. We refer to this as differential allele-specific expression (DASE) analysis. In total, we identified 341 and 260 genes (fig. 5, supplementary tables S9, S10) with DASE between highland and lowland derived F_1_ plants from the Mexican and South American continents, respectively, in at least one site:tissue by metanalysis using a 5% *lfsr* threshold. The number of genes that were significantly differentiated in ASE between highland and lowland landraces on each continent was slightly higher than the number of genes that were differentially expressed between the Mexican and South American populations on average (249, supplementary table S10) and was much higher than the number of genes that were associated with latitude on either continent (17 and 23 in the Mexican and South American populations, respectively, supplementary table S10). However, more genes showed significant changes in allele-specific expression during the approximately 1.5hr sampling window within each site:tissue (760 genes across the 3 site:tissues, supplementary table S10), which was consistent with our observations in the gene expression analysis above.

**Fig.5.**
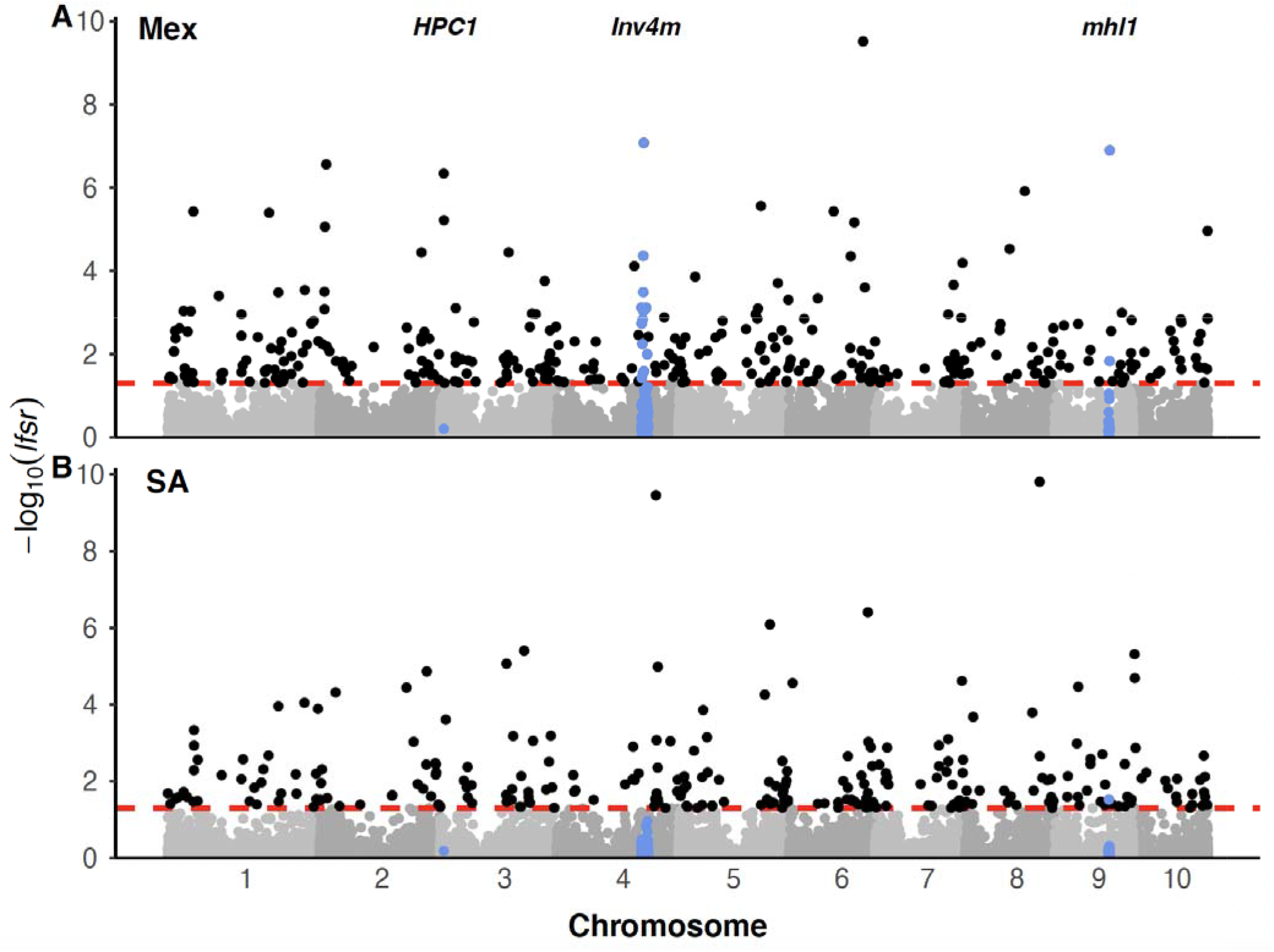
Manhattan plots showing the local false sign rate (*lfsr*) of the meta-analysis with mash for detecting differential allele-specific expression between highland and lowland landraces in the (A) Mexican and (B) South American F1 populations, expressed as −log 10 (*lfsr*). The lfsr is analogous to a false discovery rate but more stringent (Stephens, 2017). Each dot represents a gene. The dashed lines in each plot indicats the significant level at lfsr < 0.05. Blue dots highlight genes within the three prior-identified loci: HPC1, Inv4m and mhl1 and not being significant, respectively. Mex=Mexican F_1_ population, SA=South American F_1_ population.

Subsequently, we inspected the three loci with known genetic differentiation between highland and lowland landraces in Mexico: *Inv4m, mhl1* and *HPC1*. There were 13 and 2 DASE genes detected inside the *Inv4m* and *mhl1* regions in the Mexican population, but only one gene with weak evidence (*lfsr*=0.03) in the *mhl1* region in the South American population (fig. 5). This is consistent with our knowledge that the *mexicana*-to-maize introgression mainly happened in Mexican highlands (Hufford et al. 2013; Pyhäjärvi et al. 2013; Wang et al. 2017; Crow et al. 2020; Calfee et al. 2021). The differences of landrace allele-specific expression were not significant in the *HPC1* gene in either population.

Beyond the genes detected in the genomic regions that have been characterized (Hufford et al. 2013; Pyhäjärvi et al. 2013; Crow et al. 2020; Rodríguez-Zapata et al. 2021)□, the remaining genes with differentiated *cis-*regulation between highland and lowland landrace alleles had not been reported in previous studies and were distributed through all 10 chromosomes with no obvious clustering (fig. 5). We compared this list of genes (i.e., DASE), to the genes previously identified with differential gene expression (i.e., DE), between highland and lowland landraces. Of the 4,432 and 1,816 DE genes detected in the Mexican and South American populations, respectively, roughly 70% (3364 and 1235) were successfully assayed for ASE (supplementary fig. S8A). 168 and 91 genes were detected in both differential gene expression analysis and differential allele-specific analysis (supplementary fig. S8B), which account for 50% and 35% of the total numbers of DASE genes detected in the two populations.

### Convergent *cis*-regulatory evolution between the Mexican and South American populations

Among these genes that were significantly differentiated in ASE between highland and lowland populations (fig. 5, supplementary table S9), 20 were significantly associated with elevation in the F_1_ families of both continents, representing 8% of the lesser of the number of significant genes from either continent (fig. 6A, table 2). However, despite being a relatively small overlap, this is many more than expected by change (p=8.74×10^-6^). In addition, each of the 20 genes showed the same direction of changes of ASE between highland and lowland populations in both continents and the estimated highland effects of the 20 genes (r=0.93) were much more highly correlated between continents than that of all measured genes (r=0.12, fig. 6B). Therefore, we classified these 20 genes as showing convergent *cis*-regulatory evolution between the two continents.

**Fig.6.**
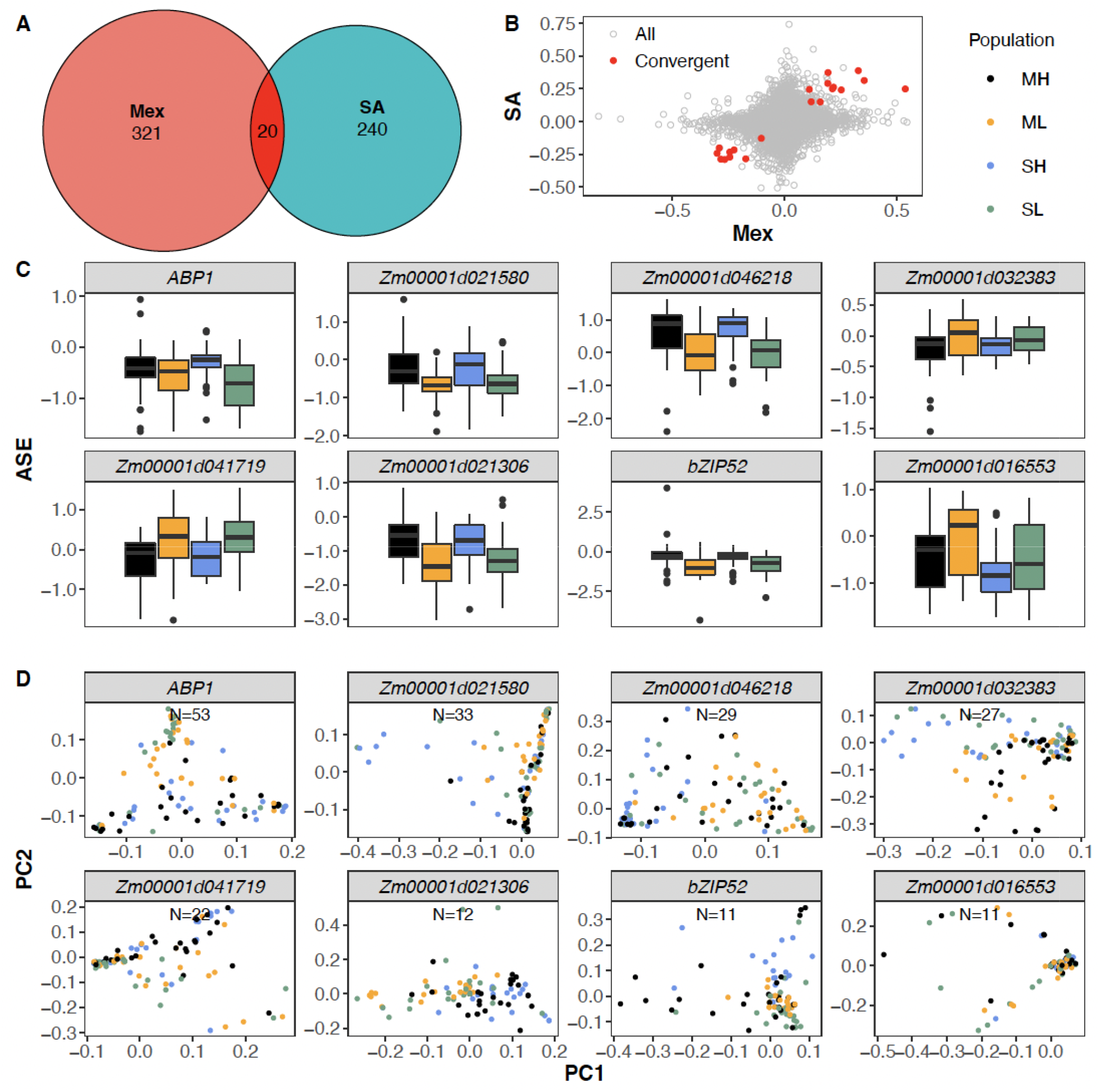
Results of allele-specific expression analyses. (A) Numbers of genes showing *cis*-regulatory divergence between highland and lowland populations from Mexico and South America and common genes detected in both continents. (B) Correlation of estimated highland effects between Mexican and South American populations for all genes measured for ASE (in gray) and a subset of 20 genes showing evidence of convergent evolution (in red). (C) ASE values of 8 of the 20 convergent genes in the Mexican Highland (ML), Mexican Lowland (ML), South American Highland (SA), and South American Lowland (SL) populations. The 8 genes were selected based on a threshold of more than 10 SNPs from the landrace parents in each of the 20 convergent genes. (D) Principal component analysis of the landraces based on SNPs called from the whole-genome sequencing data for each of 8 genes with more than 10 SNPs.

**Table 2.**
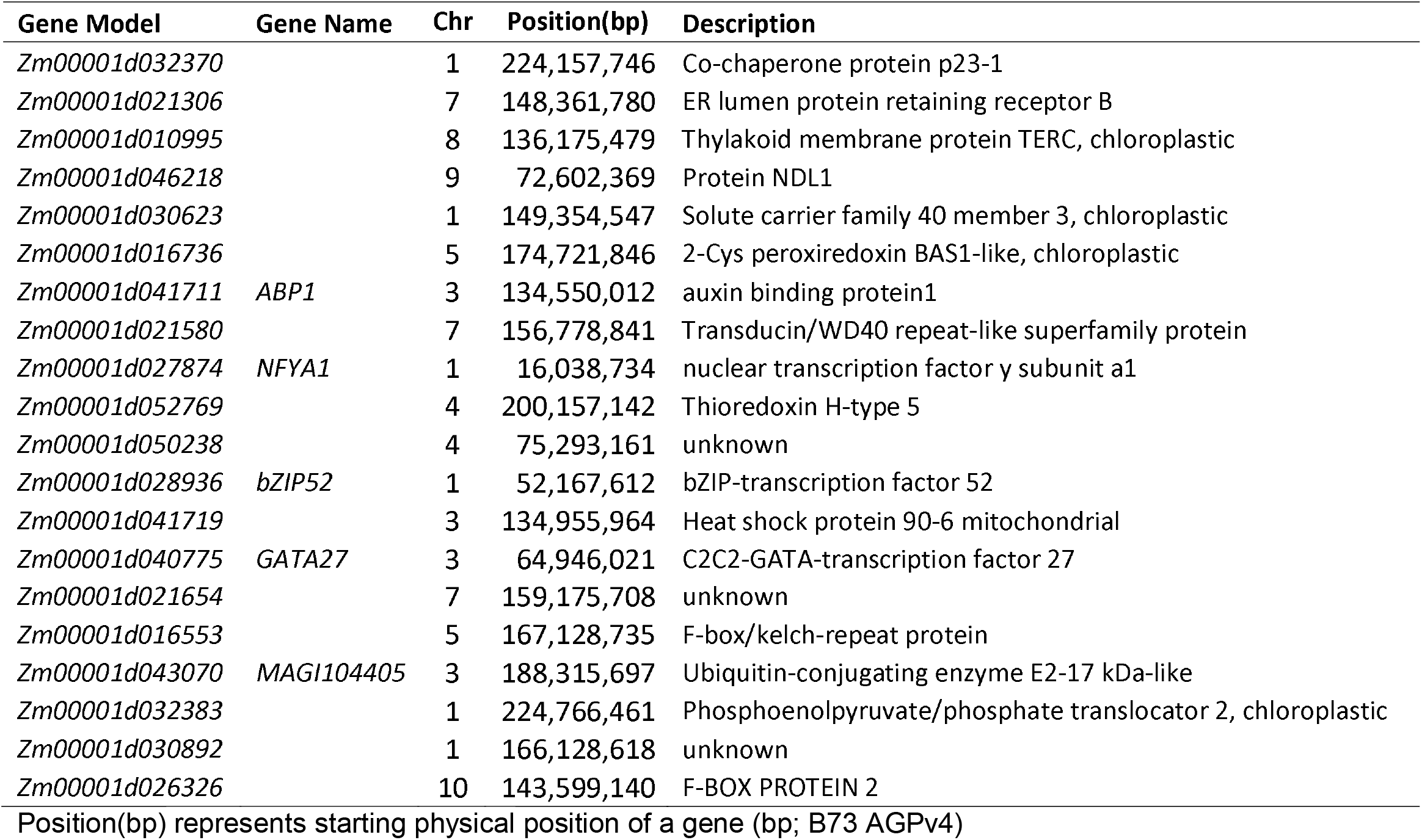
20 genes with convergent highland *cis*-regulatory evolution in both the Mexican and South American populations

| Gene Model | Gene Name | Chr | Position(bp) | Description |
| --- | --- | --- | --- | --- |
| Zm00001d032370 |  | 1 | 224,157,746 | Co-chaperone protein p23-1 |
| Zm00001d021306 |  | 7 | 148,361,780 | ER lumen protein retaining receptor B |
| Zm00001d010995 |  | 8 | 136,175,479 | Thylakoid membrane protein TERC, chloroplastic |
| Zm00001d046218 |  | 9 | 72,602,369 | Protein NDL1 |
| Zm00001d030623 |  | 1 | 149,354,547 | Solute carrier family 40 member 3, chloroplastic |
| Zm00001d016736 |  | 5 | 174,721,846 | 2-Cys peroxiredoxin BAS1-like, chloroplastic |
| Zm00001d041711 | ABP1 | 3 | 134,550,012 | auxin binding protein1 |
| Zm00001d021580 |  | 7 | 156,778,841 | Transducin/WD40 repeat-like superfamily protein |
| Zm00001d027874 | NFYA1 | 1 | 16,038,734 | nuclear transcription factor y subunit a1 |
| Zm00001d052769 |  | 4 | 200,157,142 | Thioredoxin H-type 5 |
| Zm00001d050238 |  | 4 | 75,293,161 | unknown |
| Zm00001d028936 | bZIP52 | 1 | 52,167,612 | bZIP-transcription factor 52 |
| Zm00001d041719 |  | 3 | 134,955,964 | Heat shock protein 90-6 mitochondrial |
| Zm00001d040775 | GATA27 | 3 | 64,946,021 | C2C2-GATA-transcription factor 27 |
| Zm00001d021654 |  | 7 | 159,175,708 | unknown |
| Zm00001d016553 |  | 5 | 167,128,735 | F-box/kelch-repeat protein |
| Zm00001d043070 | MAGI104405 | 3 | 188,315,697 | Ubiquitin-conjugating enzyme E2-17 kDa-like |
| Zm00001d032383 |  | 1 | 224,766,461 | Phosphoenolpyruvate/phosphate translocator 2, chloroplastic |
| Zm00001d030892 |  | 1 | 166,128,618 | unknown |
| Zm00001d026326 |  | 10 | 143,599,140 | F-BOX PROTEIN 2 |

Position(bp) represents starting physical position of a gene (bp; B73 AGPv4)

To understand the biological functions of the 20 DASE genes that were significantly associated with elevation in both continents, we searched their annotation from the Gramene database (Tello-ruiz et al. 2022)□ and their characterized function from maizeGDB (Woodhouse et al. 2021)□. 5 of them have gene names from the maizeGDB, and at least 3 of them are transcription factors (table 2). The gene *Zm00001d041711* encodes auxin binding protein 1 (ZmABP1), which binds auxin and is a receptor for a number of auxin responses (Sauer and Kleine-Vehn 2011)□. The genes *Zm00001d027874, Zm00001d028936 and Zm00001d040775* encode Nuclear transcription factor Y subunit A1 (NFYA1), bZIP transcription factor bZIP52, and GATA transcription factor GATA27, respectively. These transcription factors and transcription factor families play important roles in plant development, growth, and abiotic stress responses (Zhang et al. 2016; Guo et al. 2021; Zhang et al. 2021; Li et al. 2022)□.

For each of the 20 genes that showed consistency in both expression scales and directions in the two continents (fig. 6C, table 2), we performed a principal component analysis of the landraces based on SNPs called from the whole-genome sequencing data. We analyzed 8 genes with more than 10 SNPs each and found that landraces were separated by elevation for at least 6 genes. Highland landraces from Mexico and South America were clustered together for *ABP1, Zm00001d046218, Zm00001d041719* and *Zm00001d021306* and showed divergence for *Zm00001d021580* and *bZIP52* (fig. 6D).

### Identifying links between DASE and DE

While *cis-*regulatory variation should contribute to the total gene expression variation among samples, other sources of variation due to developmental, environmental, or *trans*-regulatory variation may dominate the gene expression variation for many genes (Liu et al. 2019). We observed generally positive correlations between log2ASE and log2Expression for most genes in each site:tissue (Supplementary fig. S9). The correlation between log2ASE and log2Expression increased when we accounted for technical factors (sampling group, order of sampling, and block), and cell type composition. However, for the majority of genes log2ASE only explained a few percent of the total expression variation.

Since several of our candidate genes for *cis-*regulatory adaptation are transcription factors, we used the MaizeGRN dataset (Zhou et al. 2020)□ which contains predicted gene regulatory networks for ~2,000 transcription factors based on co-expression results across multiple maize datasets. For each transcription factor network, we used *goseq* as described above to test whether the network was enriched for up- or down-regulated genes between highland and lowland populations. A total of 216 networks were significantly enriched in the Mexican population and 55 in the South American population in at least one site:tissue (supplementary table S11). However, we did not find any examples of these networks with transcription factors for which we observed significant divergence in *cis-*regulatory alleles in either population.

## Discussion

### Complex process of maize high-elevation adaptation

Previous studies have demonstrated substantial differences in phenotype (Anderson and Cutler 1942; Eagles and Lothrop 1994; Janzen et al. 2022)⍰ and gene expression (Kost et al. 2017; Rocío Aguilar-Rangel et al. 2017; Crow et al. 2020) between highland and lowland maize.⍰ However, the genetic architecture of regulatory variants that control these phenotypic and expression traits is still unclear and cannot be directly determined either with analyses of sequence variation or differential gene expression analysis. Differential gene expression studies cannot identify how many independent loci across the genome control expression of these genes because a single locus could plausibly affect the expression of every other gene in the genome by altering processes like cellular physiology, tissue anatomy, or organismal level development. Allele-specific expression, in contrast, as studied in the maize highland adaptation context by Aguilar-Rangel et al. (2017), is not sensitive to these *trans-* regulatory mechanisms because the two alleles of a gene are always observed in the same cellular environment. Therefore, most genes identified by Aguilar-Rangel et al. (2017) are likely controlled by distinct functional variants in *cis* to each gene⍰. However, since this study used only a single highland and a single lowland genotype, it is unclear which of the *cis-*regulatory differences they observed are common in highland populations and which may be unique to this particular lineage.

Therefore, we used population-level allele specific expression analysis, which allows us to count at least a lower-bound of the number of independent genetic loci that have diverged between highland and lowland populations. Of the 13,632 genes we successfully assayed for ASE in at least one site:tissue, 341 and 260 genes (fig. 6A) showed significantly differential allele-specific regulation between highland and lowland populations in Mexico and South America, respectively, and these genes were distributed across all 10 chromosomes with no obvious clustering (fig. 5). It is reasonable to expect more DASE genes would be detected if all the 36,207 expressed genes in maize (Hoopes et al. 2019)□ were analyzed across multiple tissues. Therefore, our results suggest a complex genetic architecture of *cis*-regulatory variants driving expression of genes for highland adaptation in maize. Furthermore, since our DASE analysis cannot detect functional variants in protein sequence or activity, for example transcription factor DNA binding affinity or other *trans*-regulatory variants, our list of candidate regulatory variants is clearly an underestimate of the total genetic architecture underlying highland adaptation. For example, recent studies have estimated that 70% or more of total expression variation in any gene is caused by *trans* effects, not *cis* effects (Liu et al. 2019)□. While some of these *trans* effects may be caused by *cis* effects on upstream genes, we have likely underestimated the number of functional variants that differ between highland and lowland maize populations.

### Evolutionary patterns of maize highland adaptation in Mexico and South America

We found a small proportion of genes for which differential gene expression or allele-specific expression were detected in both the Mexican and South American populations (fig. 2A, supplementary fig. S1). Even when assaying higher-level processes through GO categories or KEGG pathways, we found little evidence of shared patterns among the loci with gene expression divergence. Takuno et al. (2015) investigated the molecular basis of convergent adaptation in maize to highland climates in Mesoamerica and South America and found limited evidence for convergent evolution at the nucleotide level. Using high-depth resequencing data to investigate demographic change during highland adaptation, Wang et al. (2017) detected introgression from *mexicana* to maize landraces in the highlands of Mexico, Guatemala, and the southwestern USA, but found no evidence for substantial spread of *mexicana* haplotypes to South America. Consistent with these results, our analysis of two loci shown to have adaptively introgressed from *mexicana* into highland Mexicoan maize, *Inv4m* and *mhl1*, finds evidence of DASE in the Mexican population but not in the South American population (except one gene with very weak evidence detected in *mhl1*, fig. 5). Together, both our new results and previous studies suggest that the loci underlying adaptations to highlands were largely distinct and supports the model of predominantly independent evolution to the highlands in Mexican and South American maize landraces.

Nonetheless, the small but significant overlap of convergent genes detected from either differential gene expression or differential allele-specific expression in both continents suggests convergent evolution plays a non-negligible role in highland adaptation. While the genetic basis of convergence at the expression level is not clear from differential expression data alone, convergence at the *cis-*regulatory level implies functionally similar local regulatory alleles differentiating highland and lowland accessions on both continents. There are three possible mechanisms of convergence adaptation: independent mutation, shared ancestral standing variation, or spread throughout subpopulations via geneflow (Lee and Coop 2017). Of the eight genes that showed convergent *cis*-regulatory evolution between the two continents based on differential allele specific expression analysis and of which we had sufficient data from the low-coverage genome sequencing to measure local genetic relationships among samples, at least four clustered by elevation with no clear separation between Mexican highland and South American highland individuals (fig. 6D), suggesting a potential homogenization of the two highland populations through gene flow, consistent with observations of (Wang et al. 2021)⍰ where the majority of shared loci between Mexican and Andes highland landraces were due to migration. In addition, we also found at least two genes for which accessions clustered by elevation (fig. 6D), but Mexican highland and South American highland individuals clustered separated from each other. This suggests different haplotypes have arisen and/or spread independently in the two highland populations but that these two haplotypes likely have a similar biological function in each continent. However, our data cannot distinguish whether these haplotypes contain independent causal mutations, or both have captured the same variant segregating in the ancestral population. Therefore, both our results and those of Wang et al. (2021)⍰ suggest convergent evolution plays a role in maize highland adaptation, and that this adaptation likely occurred through a combination of migration and the parallel recruitment of standing and/or new mutations.

### Applications and limitations of population-level ASE analyses in evolutionary genetics and plant science

Most prior studies of ASE have been based on SNP-level allelic counts in single individuals (Rocío Aguilar-Rangel et al. 2017; Shao et al. 2019; Zhou et al. 2019). While observating ASE in an individual demonstrates that two *cis-*regulatory alleles differ functionally from each other, we cannot conclude from one individual that the populations that these individuals came from have diverged in *cis-* regulatory function until we have replicated the ASE results across multiple independently derived F_1_s. Lemmon *et* al. (2014) pioneered this approach in maize, demonstrating *cis-*regulatory divergence in many genes relative to its wild relative teosinte. Our experimental design was similar to Lemmon et al.’s (2014), except we used many more parental lines and crossed each to a common tester genotype (B73) to facilitate comparisons among all landrace alleles. As in this earlier study, we did not focus on discovering all functionally variable *cis-*regulatory alleles, but instead on identifying alleles with large changes in frequency between highland and lowland populations, as a signature of selection on gene regulation at this locus. In some cases, the divergence may represent a sweep of a particular haplotype (e.g., *Inv4m, mhl1* are candidates for this), in other cases divergence may be more polygenic even for a single gene, with an increase in frequency of multiple (potentially unrelated) haplotypes with similar *cis*-regulatory function. Detailed investigation of these alternatives will require a closer look at individual samples with higher coverage genome sequencing.

While our experimental design was optimized for discovering loci with divergent *cis-*regulatory activity between populations, it lacks power to describe the downstream effects of these loci on other traits. Since the functional alleles are necessarily in a heterozygous state in each F_1_ plant (because all landraces were crossed to a common tester), for any locus we only observe individuals that are either heterozygous or homozygous for one allele - we never observe individuals homozygous for both allelic states, and therefore cannot observe the full phenotypic effect of substituing alleles. The phenotypic differences that we observe are expected to be half of the effect we’d see if the loci were homozygous, but may be much less if the landrace allele is recessive. This likely explains why we do not see strong correlations between ASE and phenotypic traits. Even for the expression of a gene itself, *cis-* regulatory haplotypes often explain only a small percentage of the expression variation (Liu et al. 2019) due to the large number of sources of *trans-*regulatory effects. This is likely true in our study as well. We see evidence of large *trans-effects* caused by the time of day and changes in tissue composition across samples (fig. 3A,B), and after correcting for these sources of variation the correlations between ASE and gene expression do increase (supplementary fig. S9). Many of these *trans*-effects may ultimately be caused by *cis-*effects on other genes, potentially at other times or stages of development, but those effects cannot be discovered in our experiment itself. Further study of the biological roles of the *cis-*regulatory alleles we discovered here will require isolating them in other genomic backgrounds and replicating their effects in homozygous states.

Finally, while we have designed our experiment to answer questions about regulatory divergence among populations, we believe similar strategies could be used to identify gene-trait relationships relevant to hybrid breeding schemes. Hybrids dominate many key crops including maize. In such programs, candidate lines are evaluated by crossing to common testers. Experimental methods for assaying gene-level ASE as we have used here could be used for transcriptome-wide association studies (TWAS) in such hybrid populations. TWAS using ASE can pinpoint causal gene regulatory traits underlying key performance metrics, enabling further targeted gene editing work and breeding.

## Materials and Methods

### Plant materials

108 maize landraces (Supplementary table S1) from highland and lowland sites of Mexico and South America were chosen from the CIMMYT’s germplasm bank: 28 accessions from high elevation sites (> 2000 masl) and 28 accessions from low elevation sites (<1000 masl) of Mexico, and 26 accessions from high elevation sites (> 2000 masl) and 26 accessions from low elevation sites (<1000 masl) of South America. The landraces were paired latitudinally and east-west of the continental divide (Figure 1A), such that both landrace accessions of a pair collected from the same 1-degree of latitude bin and all pairwise distances between accessions were greater than 50 km. Each of the 108 maize landraces was used as a pollinator to cross with the inbred line B73 to produce 108 F_1_ families. Crosses were performed at Curtiss Farm at Iowa State University and in Columbia, Missouri, and an approximately balanced set of successful F_1_ families of each type (Highland/Lowland and Mexico/South America) were chosen from each site.

### Field experimental design and leaf sample collection

The F_1_ families were planted at two locations in Mexico: Puerto Vallarta and Metepec. Puerto Vallarta is located at 20°40’N 105°16’W and represents a lowland environment at approximately 7 masl. Over the course of the year, the temperature typically varies from 16°C to 32°C. Metepec is located at 19°14’N 99°35’W and represents a highland environment at approximately 2620 masl. Over the course of the year, the temperature typically varies from 7°C to 27°C. At each of the two locations, a randomized complete block design with two replications were used for the field trial design. The two landraces from the same latitudinal band were planted in consecutive 20 kernel rows.

Leaf tissue was sampled at the V4 developmental stage (collar of the fourth leaf became visible) from within 5 cm of the tip of the leaf blade (leaf tip) and within 5 cm of the leaf blade base (leaf base) at both locations from a randomly selected healthy-looking plant in the interior of each row. Both fields were sampled 4 hours after sunrise and all samples were taken within 90 minutes. Approximately 20 mg of tissue was sampled, placed into a 2 ml centrifuge tube, flash frozen in liquid nitrogen, and stored at −80 C until RNA extraction. Leaf tissues of the 108 landrace parents were collected, placed on ice, and transported to the laboratory where tissue was lyophilized and ground through bead beating or mortar and pestle prior to DNA isolation.

### RNA extraction, library preparation and Illumina sequencing of F_1_ hybrids

Leaf tissue was ground using stainless steel beads in a SPEX Geno/Grinder (Metuchen, NJ, USA). mRNA was extracted using oligo (dT) beads (DYNABEADS direct) to extract polyadenylated mRNA using the double-elution protocol. We prepared strand specific mRNA-seq libraries using the BrAD-seq protocol (Townsley et al. 2015)□ with random priming and 14 PCR cycles. Samples were quantified using the Quant-iTTM PicoGreen dsDNA kit, and then normalized to 1ng/ul. We multiplexed 96 samples for sequencing and sequenced each on 2-4 lanes of an Illumina HiSeq X platform generating 150 nucleotides (nt) paired-end (PE) sequences. Trimmomatic version 0.39 (Bolger et al. 2014)□ was used to remove the BrAD-seq adapters remnants and bases with an average base quality value below 15 within 4-bp sliding windows of each read. Entire reads were removed if the remaining length was shorter than 36 nt.

### Differential gene expression analysis in gene expression data

RNAseq reads of the F_1_ families were aligned to B73 AGPv4 using the STAR software version 2.7.2a (Dobin et al. 2013)□ and the STAR 2-pass method with default parameters (Engström et al. 2013)□. We counted reads at each locus using featureCounts v2.0.1(Liao et al. 2014)□ with default parameters. We filtered the raw count matrix separately for each tissue and estimated effect sizes for elevation of origin in each tissue separately, then combined evidence across three single-tissue analyses by meta-analysis to identify the union set of genes differentially expressed in at least one tissue. In detail:

First, in each single-tissue analysis, we removed F_1_ samples with fewer than 2 million mapped reads filtered genes using the *filterByExpr* function from EdgeR (Robinson et al. 2010), requiring at least 10 samples in one population-by-elevation class group to have at least 32 reads. This reduced the gene expression matrices of MetLeaftip, MetLeafbase and PvLeaftip to 18,369 genes × 160 samples, 20,401 genes × 164 samples, and 18,079 genes × 110 samples, respectively. A total of 21,599 genes were assayed in at least one site:tissue, and 16,851 genes in common among all the three tissues after filtering.

Then, for each tissue separately, we calculated normalization factors using the *calcNormFactors* function in EdgeR, normalized to log2(counts per million) and estimated weighting factors with voom (Law et al. 2014). To perform voom processing, for each tissue, we specified a linear model accounting for Block (in Metepec samples only), the sampling team (3 teams sampled tissue in parallel), sampling time (expressed as a cubic polynomial of the order in the field, separately for each of the 3 sampling teams), the interaction of Population (Mexico or South America) and Elevation class (Highland or Lowland parental landrace), and the interaction of Population and Latitude of the parental landrace.

Next, we re-fitted the linear model described above using lmFit in limma (Ritchie et al. 2015) taking the precision weights estimated by voom into account. We used the eBayes function to perform empirical Bayes moderation of the t-statistics. We extracted the estimated average difference in log2(counts per million) between highland and lowland-derived F_1_s for each population separately from fit$coefficients and the standard errors of these estimates as sqrt(fit$s2.post) * fit$stdev.unscaled.

Finally, we performed a meta-analysis of the elevation effects of each gene across three tissues, accounting for correlations of measurements among conditions using the multi-variate adaptive shrinkage (mash) method implemented in mashr package 0.2.50 (Urbut et al. 2019)□⍰ on the estimated effect sizes and standard errors calculated above. This produced a union set of genes with evidence of a difference in the average expression between highland and lowland F_1_s in any condition. We used the 21,599 genes with estimated elevation effects in at least one site:tissue for the meta-analysis, setting input effect sizes and output results to *NA* for genes not assayed in a particular site:tissue. We ran mashr with the *mash_estimate_corr_em* to estimate a residual correlation matrix, passing both the canonical covariance matrices (*cov_canonical*) and data-driven covariance matrices (*cov_ed*, with inputs from *cov_pca* pasted on the genes significant at a *lfsr* of 0.05 in at least one condition).

### Gene set analysis in gene expression data

We ran gene set enrichment analyses on gene lists discovered by the meta-analysis across tissues, separately for the Mexican and South American populations, using the *goseq* function of the *goseq R* package (Young et al. 2010)□. We began with a list of 12,035 Gene Ontolog (GO) categories (Wimalanathan et al. 2018)□, 137 KEGG pathways (Kanehisa et al. 2021), and 556 CornCyc pathways (Hawkins et al. 2021)□, and then filtered for categories with between 10 and 1000 assayed genes in a particular site:tissue. We ran the enrichment analyses separately for up- and down-regulated genes selected with by *lfsr* < 0.05 in each site:tissue. We accounted for biased probabilities of detection as a function of expression and gene length using the *nullp* function with *bias.data* set to the log of the average counts per gene across all samples in that site:tissue, including only genes that passed the expression filter described above.

We assessed convergence in each site:tissue at the gene level by selecting genes with *lfsr < 0.05* for effects of elevation separately in the Mexican and South American populations and filtering for genes where the Posterior Mean effect size estimate had the same size in both populations. We assessed convergence at the gene set level based on Benjamini-Hochberg adjusted p-values < 0.05 in the test of either up-regulated or down-regulated genes for both populations.

### Assessment of cell composition variation among samples

We used single-cell expression data from Bezrutczyk et al. (2021) to estimate cell composition in each sample. This dataset included 200-900 marker genes with enriched expression in 7 cell types (5 classified as mesophyll and 2 as bundle sheath). We calculated a projection score for each of our samples against each of the 7 cell as the weighted sum of mean-centered expression of the marker genes (weighted by the *avg_log2FC* in the specific cell population in the reference dataset). This is closely related to the OLS method for estimating cell type proportions in single-cell expression data (Avila Cobos et al. 2020)□, but less restrictive because we do not assume that all cell populations in our samples are represented in the reference dataset. We summarized variation in cell type composition across samples using a principal components analysis of the 7 projection scores.

To assess the reliability of the projection scores we re-calculated the scores 200 times after randomly assigning the marker gene identities to random expressed genes and measuring the total variation explained by the real or permuted scores across samples.

We assessed whether the projection scores representing cell composition variation could account for some of the differential expression observed between highland and lowland-derived F_1_s by including the 7 projection scores as additional covariates in the design matrices for the differential expression analyses derived above.

### Whole-genome sequencing and variant identification from the landrace parents

Since variant calling from RNAseq libraries is notoriously difficult due to: (i) allelic imbalance, since most variant callers assume the true frequency of each allele is 50%, (ii) highly variable sequencing coverage across loci, negating depth filters from variant calling software, and (iii) mapping difficulties due to spliced reads, we used low-coverage whole-genome sequencing data of the landrace parents to identify a set of high-confidence genic SNPs to use for ASE quantification.

DNA was extracted from parental landrace leaf tissue using the CTAB method. The tissue was collected from the same male plant used to produce the F_1_s that were used for RNA sequencing. Sample concentrations were quantified using Qubit (Life Technologies), and 1ug of DNA was fragmented using a bioruptor (Diagenode) with cycles of 30 seconds on, 30 seconds off. Fragments of DNA were then prepared for Illumina sequencing. (1) DNA fragments were repaired with the End-Repair enzyme mix (New England Biolabs). (2) A deoxyadenosine triphosphate was added at each 3’end with the Klenow fragment (New England Biolabs), and (3) Illumina Truseq adapters (Affymetrix) were added with the Quick ligase kit (New England Biolabs). Between each enzymatic step, DNA was washed with sera-mags speed beads (Fisher Scientific). Finally, samples were multiplexed using Illumina compatible adapters with inline barcodes and libraries were sequenced with Illumina HiSeq X platform generating 150 nucleotides (nt) paired-end (PE) sequences, resulting in an average of 9,862,996 properly paried reads/library, corresponding to an average of ~1.2x coverage. Reads were aligned to version 4 of the B73 reference genome (Jiao et al. 2017)□ with BWA-MEM version 0.7.17 (Li and Durbin 2009)□. High-confidence SNPs between any landrace and B73 were identified with Analysis of Next Generation Sequencing Data (ANGSD) version 0.931-2 (Korneliussen et al. 2014)□ using the following parameters: angsd -GL 2 -P 20 -uniqueOnly 1 -remove_bads 1 - only_proper_pairs 1 -trim 0 -C 50 -minMapQ 20 -mminQ 20 -SNP_pval 1e-6 -doMaf 2 -doMajorMinor 4 -doSaf 1. SNPs outside of annotated exons in the B73 genome were excluded.

Since the landrace parents were outbred, their genomes are heterozygous and the ~1x whole-genome sequencing (WGS) reads will likely not detect ~50% of the SNPs caried by each parent and passed on to the F_1_ individuals. Given the size of the maize genome, achieving sufficiently high coverage for each individual for comprehensive SNP discovery would have been prohibitively expensive. However, SNPs relative to the reference genome (B73 AGPv4) that are relatively common in the population (e.g. > 2% frequency) are likely to be sequenced by multiple reads across all 108 WGS libraries. This includes a large number of SNPs where the B73 allele is rare which will be observed in nearly every landrace. In total, we identified 53,891,495 high-confidence SNPs in exonic regions across the 108 landraces, providing a large set of candidate SNPs to test for ASE in the RNAseq data.

### Per-sample detection of ASE-tagging SNPs without biasing ASE ratios

While the WGS-derived SNPs are likely real in the whole population, only SNPs that are heterozygous in a particular F_1_ individual are useful for ASE quantification. Including the same set of fixed loci in ASE counts across samples will severely bias allelic read counts for a gene because all reads from both alleles will be assigned to the same allele. We therefore used the RNAseq data to genotype each F_1_ individual at all WGS-derived SNPs.

Using WGS-derived SNPs alleviates the issue of confident SNP detection, but genotyping using RNAseq data for ASE applications still presents challenges:

i. When a small number of reads cover a SNP (e.g. when in a low-expressed gene) one allele will frequently drop-out due to sampling error even if there is no actual allelic imbalance. In our experimental design, we know that every locus contains at least one copy of the B73 allele (since B73 was the female parent). While loci where only the landrace allele was observed are almost certainly heterozygous and therefore informative for ASE, keeping these loci would bias the genes estimated ASE ratio towards the landrace allele, because the opposite loci (where only the B73 allele is detected) would be dismissed as apparently homozygous. We therefore kept only SNPs where both the B73 and the landrace allele were observed to prevent biased ASE ratios.
ii. When a large number of reads covers a SNP (e.g. when in a high-expressed gene), the low rate of sequencing errors present in Illumina data can generate false-positive heterozygous calls. Including these loci in the ASE analysis will severely bias ASE ratios towards the B73 allele (because most sequencing errors will be away from the reference and therefore look like low-expressed non-B73 alleles.
iii. Mismatches relative to the reference can cause ambiguous or incorrect read-mapping, biasing ASE ratios. We used the WASP algorithm (Van De Geijn et al. 2015)□□ implemented in the STAR software version 2.7.2a to identify reliably mapped reads. WASP uses an allele swapping and RNA-seq remapping strategy to filter out reads with mapping biases□, and the STAR-WASP algorithm assigns a multi-locus genotype to each individual read for all SNPs it overlaps.

RNAseq reads of the F_1_ families were aligned to B73 AGPv4 using the STAR software version 2.7.2a and the STAR 2-pass method was used with default parameters□. For each F_1_ sample separately, alleles were counted at WGS-derived loci using ASEReadCounter from GATK version 4.0.11.0. To minimize the impact of the above issues on downstream ASE analyses, we kept only SNPs for each sample where both alleles were detected, the total number of reads covering the SNP was at least 10, and the absolute value of the log2ASE ratio: log2(ALT)-log2(REF) was less than 2. We applied these filters to each SNP in each RNAseq sample.

### Identifying regions of IBD between plants from the same F_1_ family

We used the heterozygous SNP calls from each RNAseq sample to identify regions of IBD between the three plants per F_1_ family (two plants from two blocks in Metepec and one plant from Puerta Vallarta). For each F_1_ family, we compared RNAseq samples of two tissues from the same plant in Metepec and of two plants from two blocks in Metepec/Puerta Vallarta for the same tissue. For each pair of RNAseq samples, we divided each chromosome into 20 blocks with equal numbers of SNPs from the WGS data, and in each bin counted the number of heterozygous sites identified in common between the two samples. We then divided this number by the minimum number of heterozygous sites identified in each sample separately. This percentage of common sites was generally bimodal across bins, reflecting the inheritance of the two paternal alleles in the sibling plants. We fit a gaussian mixture distribution to these percentages for each sample with k=2 using the normalmixEM function from the mixtools package (Benaglia et al. 2009)□ to classify each bin into either IBD (if the posterior probability of the bin being in the higher-probability class was > 90%), not-IBD (posterior-probability < 10%), or ambiguous.

### Gene-level allelic read counts for F_1_ samples

While SNP-level allelic expression counts can document allelic imbalance in a single sample, to identify genes with common allelic imbalance at the population level we combined the information across SNPs in the same gene into a single ASE ratio per gene per sample. Gene-level ASE ratios should be more robust because they are based on more total reads, and in a population sample SNP-level ASE ratios cannot reliably be compared across individuals because many SNPs are individual-specific.

To combine SNP-level allelic expression counts into gene-level allelic expression we used the WASP algorithm (Van De Geijn et al. 2015)□□ implemented in STAR-WASP (Dobin et al. 2013)□. Therefore, we extracted reads that were assigned either REF or ALT genotypes at all overlapping loci into separate BAM files, and then counted the reads overlapping each gene feature in each BAM file using featureCounts v2.0.1 (Liao et al. 2014)□. These gene counts are the allelic expressions of the maternal and paternal alleles of each gene, respectively.

### Differential allele-specific expression analysis

Using the gene-level allelic read counts, we analyzed the average difference in landrace allele-specific expression (relative to B73 allele-specific expression) between F_1_s derived from highland and lowland landraces. We modeled this landrace elevation effect separately for three tissues: the leaf tip and leaf base tissues from the Metepec field (MetLeaftip, MetLeafbase), and the leaf tip samples from the Puerta Vallarta field (PvLeaftip). We then performed a meta-analysis across three tissues to identify the set of genes with divergent allelic expression between highland and lowland F_1_s in any condition.

First, in each single-tissue analysis, we removed F_1_ samples with fewer than 2 million mapped reads and genes in which fewer than 10 samples had at least 32 ASE-informative reads *in each of the 4 populations*. This stronger filter was necessary for the ASE analysis because genes with few reads are informative for total expression analyses (*i.e*. low expressed), but uninformative for ASE. For each gene in each F_1_ sample, we calculated the log2ASE ratio as log2(landrace counts) – log2(B73 counts), where landrace and B73 are actually paternal and maternal alleles, respectively. This resulted in datasets of size: 10,886 genes × 160 samples for MetLeaftip, 12,747 genes × 164 samples for MetLeafbase, and 9178 genes × 110 samples for PvLeaftip. A total of 13,632 genes were assayed in at least one site:tissue, and 8,605 genes were in common among all the three tissues after filtering.

We expected that the precision of these log2ASE ratios would vary strongly among genes and samples due to the expression of each gene, the number of informative SNPs, and the sequencing depth of each sample. This heteroskedasticity would reduce the efficiency of standard tests for differential expression (similarly to the effect of counting variance on total expression in RNAseq samples). We therefore developed an adaptation of the voom algorithm for modeling the expected variance of each datapoint. For each tissue, we specified the same linear model accounting for Block, sampling group, order in the field, the interaction of Population and Elevation class, and the interaction of Population and Latitude of the parental landrace as described above in the total expression analysis. We used the lmFit function in limma version 3.42.2 (Ritchie et al. 2015)□□ to fit this model to the log2ASE ratios of each gene and extracted the estimate of the residual standard deviation of each gene. In this step, all genes with zero counts from either allele were set to missing (given zero weights) because a zero log2ASE ratios implies equal allelic expression while zero counts is a complete lack of information about the actual allelic ratio. Next, we used the lowess function to fit a smoothed trend to the square root of residual standard deviations extracted above as a function of an average normalized total counts of each gene (in log2 scale). Finally, we used this trend line to predict the variance of each observation in the data matrix as a function of the total read count (landrace + B73) of that gene in that sample.

Next, we re-fitted the linear model above using lmFit, this time including the inverse of the estimated variance matrix as precision weights, again setting the weights of points with zero total counts to zero. We used the eBayes function to perform empirical Bayes moderation of the t-statistics. We extracted the estimated average difference in log2ASE between highland and lowland-derived F_1_s for each population separately from fit$coefficients and the standard errors of these estimates as sqrt(fit$s2.post) * fit$stdev.unscaled.

Finally, based on the observed effect sizes and corresponding standard errors of each gene of three single-tissue analyses, we performed a meta-analysis using mashr (Urbut et al. 2019)□□ to identify a union set of genes with evidence of a difference in the average landrace allele-specific expression between highland and lowland F_1_s in any condition following the same procedure of total expression analysis. In this analysis, the *mash* results suggested the correlation in true effect sizes was close to 1 across all three site:tissues. We therefore used the overall *lfsr* across all three site:tissues as a measure of signficance, and did not break results down by site:tissue.

## Supporting information

supplemental results

## Supplementary Material

All supplementary figures, tables, results and text have been included in the supplementary files.

## Acknowledgements

This study was supported by the National Science Foundation (grant number 1546719). The authors thank Dr. Graham McVicker at The Salk Institute, Dr. Alexander Dobin at Cold Spring Harbor Laboratory and Mr. Arya Massarat at Harvey Mudd College for their valuable advice on building the allelic read counts pipeline. The authors also thank Dr. Garrett Janzen for his valuable advice and discussion on analysis of Mexican and South American maize landrace populations.

## Data Availability

The pipeline and custom scripts utilized in this paper are documented in the following GitHub repository:https://github.com/hh622/Maize_Highland_Adaptation_allele_specific_expression. The RNA sequencing (PRJNA796614) and the whole genome sequencing (PRJNA799784) raw reads have been deposited in NCBI SRA.

