## supplemental results for "Allele-specific expression reveals multiple paths to highland adaptation in maize"

### Supplementary Results

#### Three validation analyses to assess the reliability of our ASE pipeline

To assess how well our pipeline can control allelic mapping bias, we examined distributions of ASE ratios (in log2 scale) for each F_1_ sample. While landrace alleles may be higher or lower expressed than B73 allele for individual genes, we expected that the true distribution of log2ASE ratios across all gene for each sample should be centered around 0. A bias towards observing more reads mapped to B73 alleles is likely an artifact of mapping bias. We found that the distribution of log2ASE ratios was approximately symmetric for nearly every sample; matched upper- and lower- quantiles were nearly identical in absolute values with only slight bias to the reference alleles for larger |log2ASE| values (supplementary fig. S5A). In contrast, we found that without selecting high-quality SNPs led to the mapped allelic reads biased to the reference allele (supplementary fig. S5B).

To assess whether the observed distribution of log2ASE ratios per sample was greater than expected due to pure allele-counting error, we first stratified the ASE from our real data (hereafter real ASE) by the average expression level of the genes (quantile 1= lowest expressed, 10=highest; genes with fewer total counts will have greater sampling variation in log2ASE); and then, for each gene, we simulated ASE by sampling from a binomial distribution given its total count. By comparing distributions of real ASE against simulated ASE for groups with low, medium and high average expression levels (supplementary fig. S6), we found that our real data had much more variation in ASE than expected by chance across genes with different expression levels; variation of simulated ASE decreased with more counts, but the real ASE variation almost had no changes as expression level increased.

To further assess the reliability of ASE measured by our analysis pipeline, we compared ASE measured between tissues from same plants and between two individuals of the same F_1_ family from the same tissue. We found that the correlation of ASE between samples collected from two different individuals from the same F_1_ family was very high for genes in genomic regions where the two individuals shared the same haplotype (i.e., identical by descent, IBD), especially for genes with high numbers of informative RNAseq reads, but much lower for genes in genomic regions where the two individuals did not share the same haplotype. When we compared ASE values between the two tissues (leaf tip and leaf base) on the same plant, the correlation was lower than when we compared the same tissue across individuals in IBD regions (supplementary fig. S7).
