## supplementary text for "Allele-specific expression reveals multiple paths to highland adaptation in maize"

### Major challenges for ASE detection across individuals at the gene level when only RNAseq data is available and how we solved this challenges in our study

We used ASE (defined as ratio of landrace allelic count to B73 allelic count in log2 scale) to scan the genome for genes that have undergone adaptive divergence in the *cis*-control of gene expression between highland and lowland landrances. There are at least three major challenges for ASE detection across individuals at gene level when only RNAseq data is available. First, *cis*-regulatory variants are generally not contained within the mRNA molecules, and therefore cannot be observed with RNAseq data alone. Second, ASE is based on counting reads that overlap with exonic heterozygous SNPs (hetSNPs), however, phase information of exonic hetSNPs within a gene is usually unknown. Third, even when *cis*-regulatory variants and haplotypes of exonic hetSNPs are known, the linkage disequilibrium between them across multiple individuals are unknown. Existing studies for ASE detection are either in single individuals (Rocío Aguilar-Rangel et al. 2017; Shao et al. 2019; Zhou et al. 2019, but see Lemmon et al. 2014 who used 29 F_1_s from different maize and teosinte parents to study the genetics of maize domestication) or across individuals based on a pseudo phasing procedure in which a ‘majority voting’ approach is used to assign allelic read counts at each SNP site within each gene to the major haplotype that was defined as the haplotype has higher expression than the other haplotype (Mayba et al. 2014; Fan et al. 2020).

These challenges have been resolved with an appropriate genetic design in our study. We crossed each of the 108 maize landraces as pollinator (paternal parent) with a common maize inbred line B73 (maternal parent) to generate 108 F_1_ hybrids, such that individual seeds of the 108 hybrids carry a common gene allele from B73 and a second gene allele from landrace at each gene locus. When mapping RNAseq reads of F_1_s onto the B73 genome (AGPv4), origins (landrace or B73) of RNAseq reads can be determined according to haplotypes of all hetSNPs that they overlap under our genetic design. Further, for each F_1_ sample at each gene locus, we know the allelic count of the landrace allele was regulated by landrace *cis*-regulatory allele and the allelic count of B73 allele was regulated by B73 *cis*-regulatory allele. Furthermore, although every landrace possibly has its own *cis*-regulatory allele that comes from a population with different coalescent history, for loci underlying highland adaptation, different *cis*-regulatory haplotypes within a population (either highland or lowland) possibly have similar biological functions. By testing average difference in ASE values for each gene between highland and lowland samples, we could identify *cis*-regulatory divergence at the population level between highland and lowland alleles in the Mexican and South American populations.
