## supplementary fig. S1 for "Allele-specific expression reveals multiple paths to highland adaptation in maize"

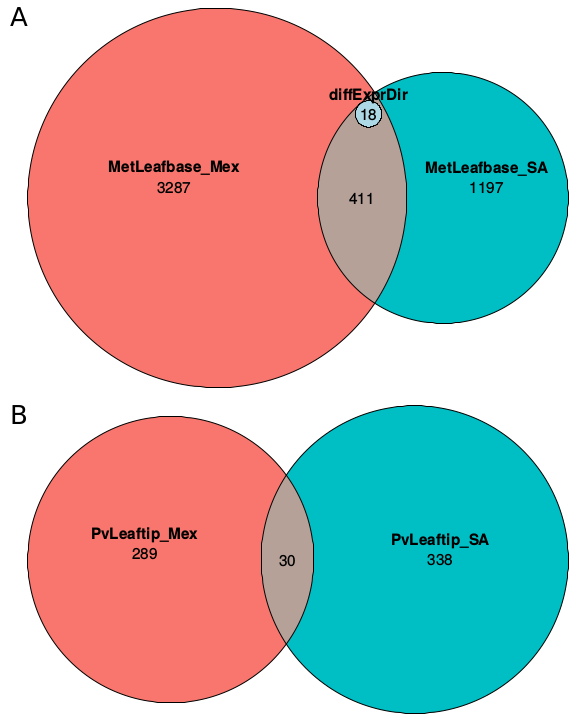

**Supplementary Figure 1.** Numbers of differentially expressed genes between highland and lowland populations from Mexico (red) and South America (blue) and common genes detected in both continents. Analyses of F1 samples from the two populations were done separately post-normalization. (A) MetLeafbase and (B) PvLeaftip. The small inset in the overlapping region of figure A, shows genes significant in both populations, but with opposite directions of expression change.
