## supplementary fig. S2 for "Allele-specific expression reveals multiple paths to highland adaptation in maize"

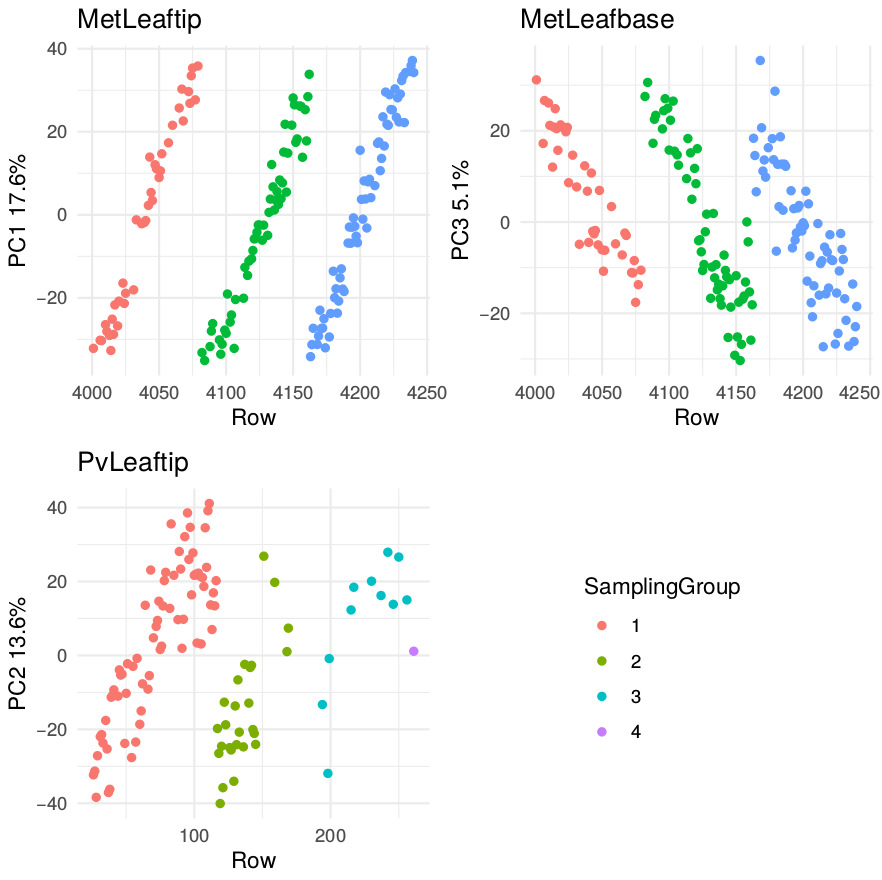

**Supplementary Figure 2.** Associations between principal components of gene expression and sampling times within the 1.5 hr sampling window for the three site:tissues. Each dot represents one F_1_ sample. Points are arranged on the x-axis according to the field row (rows snaked back and forth across the field in a set of 5 (Metepec) or 10 (Puerta Vallarta) ranges). Colors represent the 3-4 sampling teams that sampled plants in parallel during the sampling window. For this figure we chose the PC with the strongest association with Row (i.e., order of sampling) to demonstrate that this was an important component of expression variation.

level.
