## supplementary fig. S3 for "Allele-specific expression reveals multiple paths to highland adaptation in maize"

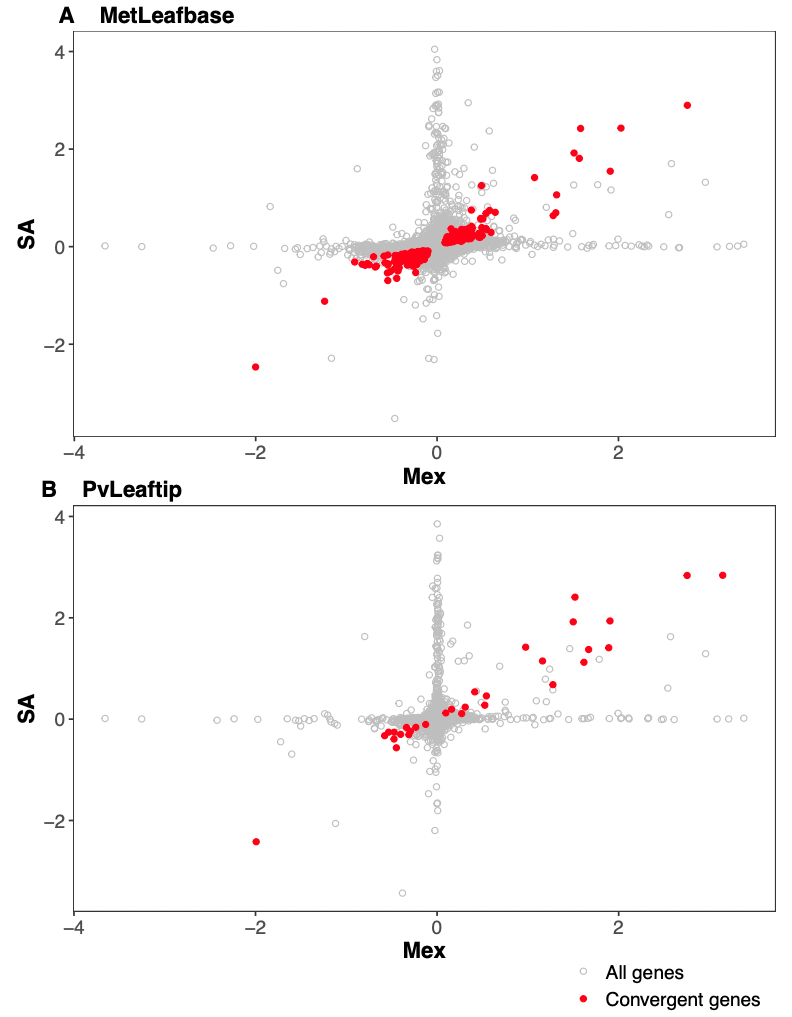

**Supplementary Figure 3.** Correlation of posterior mean highland effects between Mexican and South American population for all genes measured for gene expression (in gray) and a subset of genes showing evidence of convergent evolution (in red) in (A) MetLeafbase and (B) PvLeaftip.
