## supplementary fig. S4 for "Allele-specific expression reveals multiple paths to highland adaptation in maize"

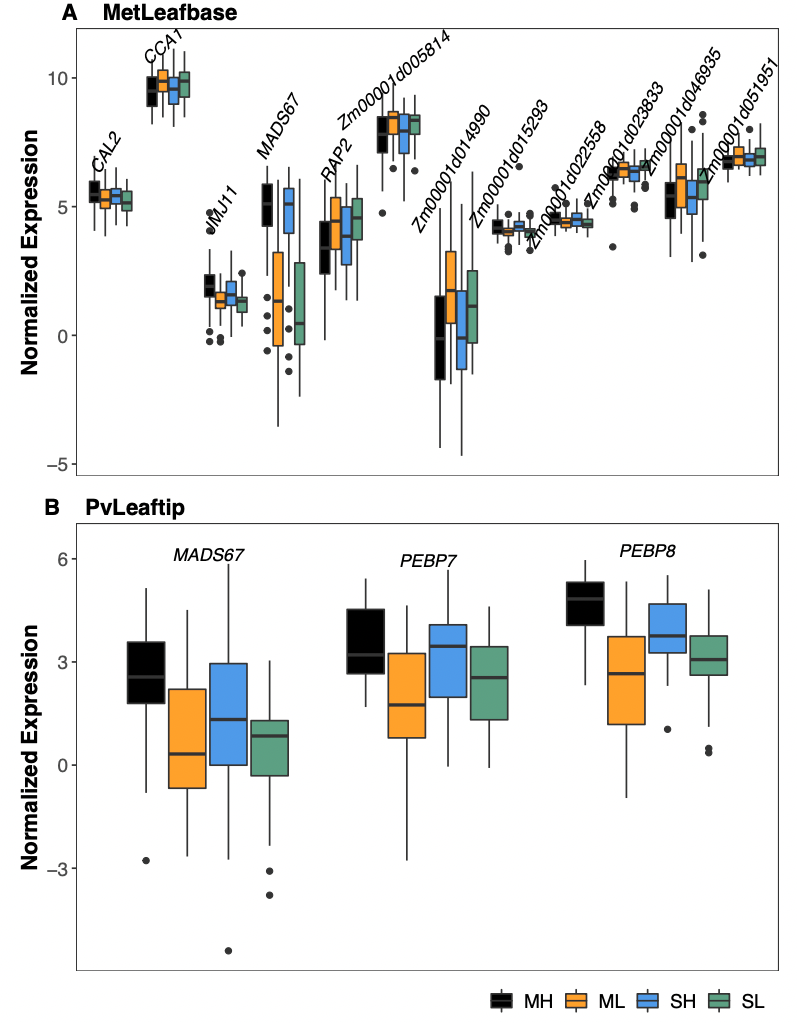

**Supplementary Figure 4** Expression of flowering-related genes in the Mexican Highland (ML), Mexican Lowland (ML), South American Highland (SA), and South American Lowland (SL) populations in A) MetLeafbase and (B) PvLeaftip. These flowering-related genes are identified by looking for overlapping between the convergent genes and maize flowering time candidate genes aggregated by Li et al. (2016) and Swarts et al. (2016).
