## supplementary fig. S5 for "Allele-specific expression reveals multiple paths to highland adaptation in maize"

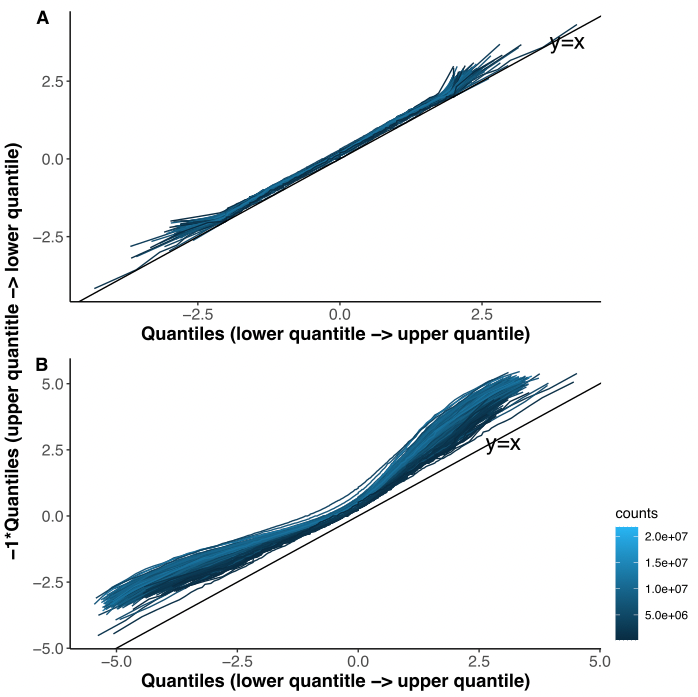

**Supplementary Figure 5.** Filtering for SNPs with at least one count from both alleles reduces reference bias in ASE ratios. Q-Q plots of quantiles of log2ASE ratios across all genes (x axis) against -1 * the quantiles of log2ASE ratios in reverse order (y axis). Reference bias (tendency of higher counts for the reference allele) shows up as up-ward shifts relative to the y=x line. (A) filtered: heterozygous SNPs for each sample, and (B) unfiltered: all heterozygous SNPs for each sample where both alleles were detected, the total number of reads overlapping the SNP was at least 10, and the absolute value of the log2ASE ratio (i.e, log2(Landrace/B73)) was no larger than 2.
