## supplementary fig. S6 for "Allele-specific expression reveals multiple paths to highland adaptation in maize"

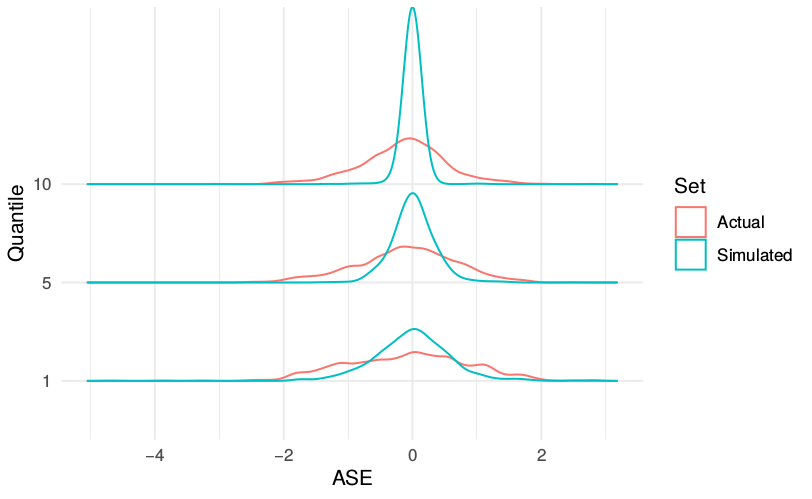
**Supplementary Figure 6.** Distributions of ASE from real data (real ASE, in blue) and from simulated data (in red) for a random sample. We first stratified the real ASE by the average expression level of the genes (quantile 1= lowest expressed, 10=highest); and then, for each gene, we simulated ASE by sampling from a binomial distribution given its total count.
