## supplementary fig. S7 for "Allele-specific expression reveals multiple paths to highland adaptation in maize"

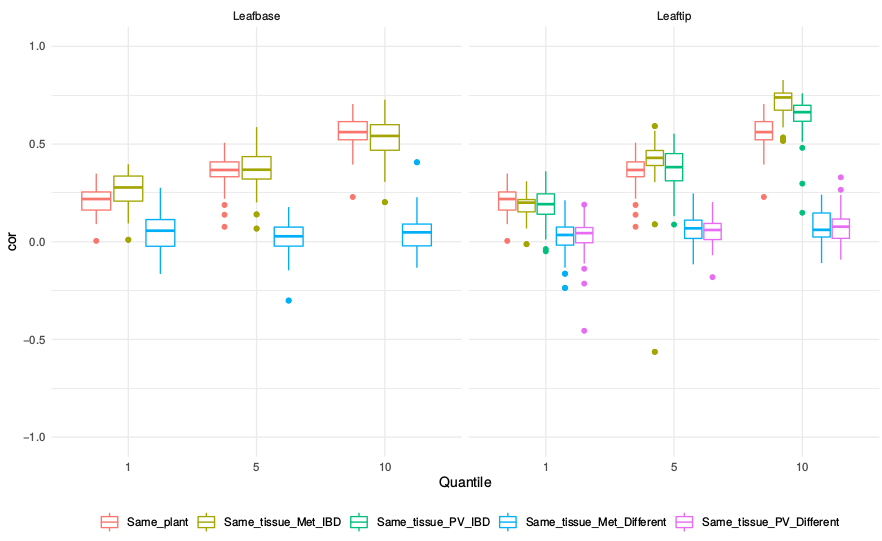
**Supplementary Figure 7.** Correlation of ASE estimates across samples. Genes are stratified by deciles of expression level (1=lowest, 10=highest), and correlations are reported comparing ASE between different tissues collected from same plant (red), between two individuals of the same F_1_ family from the same tissue for genes in genomic regions where the two individuals shared the same haplotype (IBD) (yellow = both plants in Metepec, green = one plant in Metepec, one in Puerta Vallarta) or did not share the same haplotype (blue/purple).
