## supplementary fig. S8 for "Allele-specific expression reveals multiple paths to highland adaptation in maize"

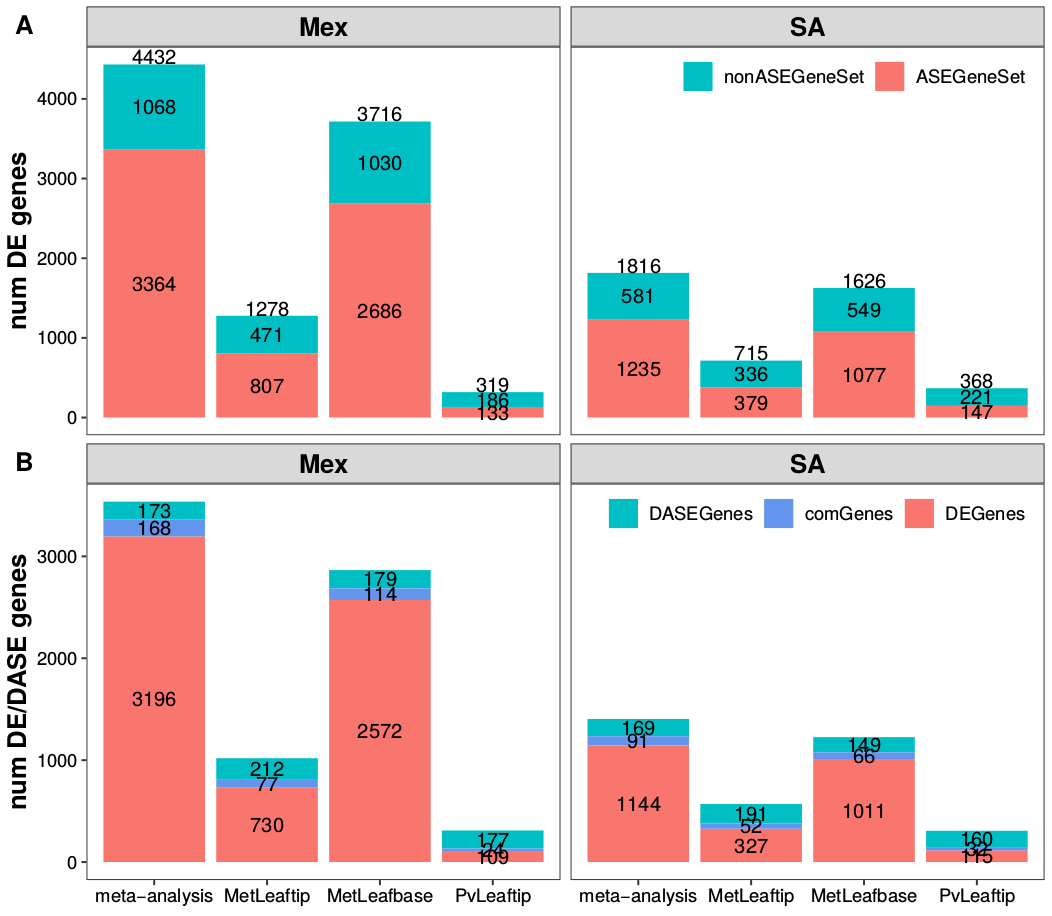

**Supplementary Figure 8.** Comparison of the number of genes detected in the differential gene expression (DGE) and differential allele specific expression (DASE) analyses. (A) Numbers of differentially expressed (DE) genes detected in single-tissue analysis and in a meta-analysis based on the same set of genes for DASE analysis (colored in red) and additional genes that were assayed for gene expression but not for ASE (colored in blue). (B) Comparison of DE genes and DASE genes detected in single tissue analysis and in a meta-analysis based on the same set of assayed genes. The numbers of common genes detected between DASE and DE analyses were highlighted in light blue.
