## supplementary fig. S9 for "Allele-specific expression reveals multiple paths to highland adaptation in maize"

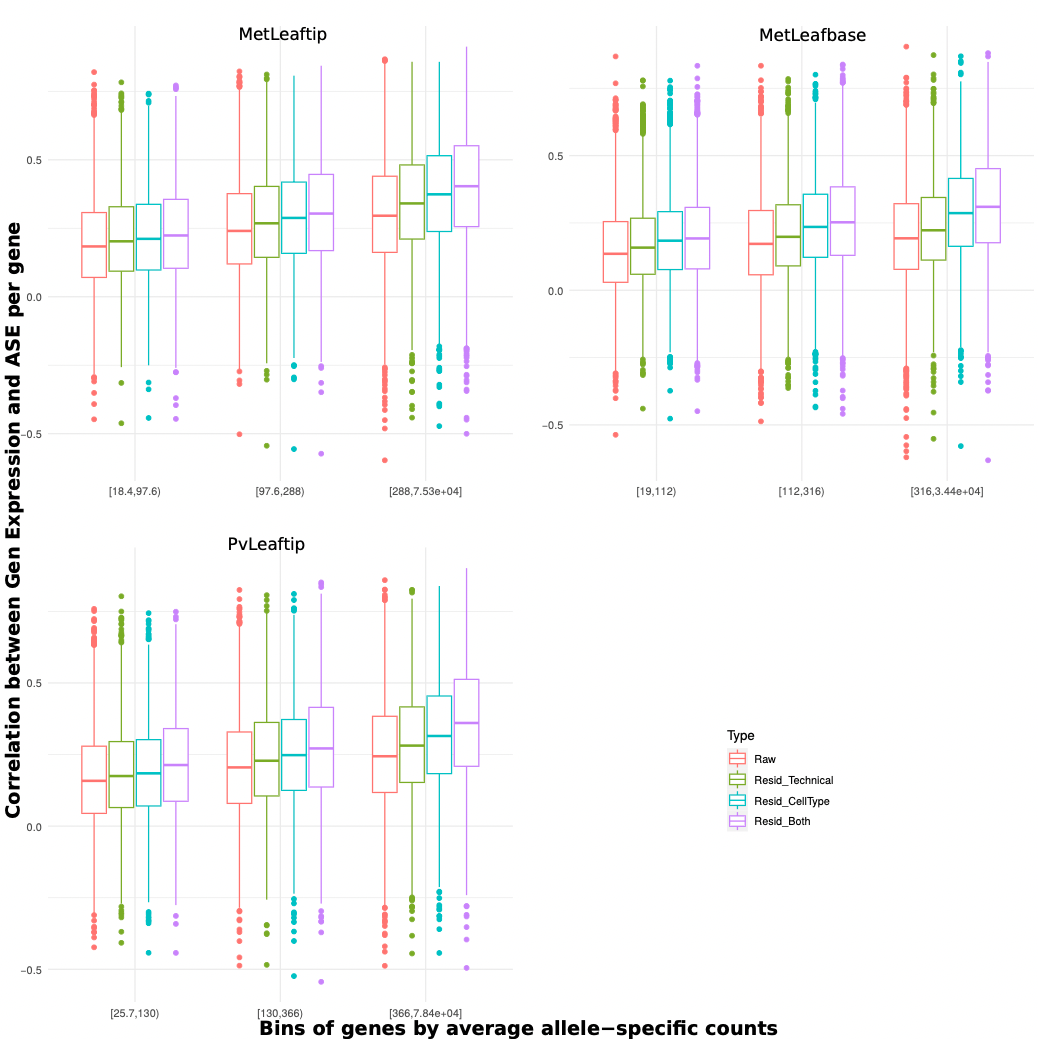

**Supplementary Figure 9.** Correlations between log2 scale ASE and log2 scale gene expression for common genes assayed for gene expression and ASE in each site:tissue. Boxplots show the distributions of correlations across genes. Genes were stratified by terciles of expression level.
